# Histone H2Bub1 deubiquitylation is essential for mouse development, but does not regulate global RNA polymerase II transcription

**DOI:** 10.1101/2021.02.23.432458

**Authors:** Fang Wang, Farrah El-Saafin, Tao Ye, Matthieu Stierle, Luc Negroni, Matej Durik, Veronique Fischer, Didier Devys, Stéphane D. Vincent, László Tora

## Abstract

Co-activator complexes dynamically deposit post-translational modifications (PTMs) on histones, or remove them, to regulate chromatin accessibility and/or to create/erase docking surfaces for proteins that recognize histone PTMs. SAGA (Spt-Ada-Gcn5 Acetyltransferase) is an evolutionary conserved multisubunit co-activator complex with modular organization. The deubiquitylation module (DUB) of mammalian SAGA complex is composed of the ubiquitin-specific protease 22 (USP22) and three adaptor proteins, ATXN7, ATXN7L3 and ENY2, which are all needed for the full activity of the USP22 enzyme to remove monoubiquitin (ub1) from histone H2B. Two additional USP22-related ubiquitin hydrolases (called USP27X or USP51) have been described to form alternative DUBs with ATXN7L3 and ENY2, which can also deubiquitylate H2Bub1. Here we report that USP22 and ATXN7L3 are essential for normal embryonic development of mice, however their requirements are not identical during this process, as *Atxn7l3*^−/−^ embryos show developmental delay already at embryonic day (E) 7.5, while *Usp22*^−/−^ embryos are normal at this stage, but die at E14.5. Global histone H2Bub1 levels were only slightly affected in *Usp22* null embryos, in contrast H2Bub1 levels were strongly increased in *Atxn7l3* null embryos and derived cell lines. Our transcriptomic analyses carried out from wild type and *Atxn7l3*^−/−^ mouse embryonic stem cells (mESCs), or primary mouse embryonic fibroblasts (MEFs) suggest that the ATXN7L3-related DUB activity regulates only a subset of genes in both cell types. However, the gene sets and the extent of their deregulation were different in mESCs and MEFs. Interestingly, the strong increase of H2Bub1 levels observed in the *Atxn7l3*^−/−^ mESCs, or *Atxn7l3*^−/−^ MEFs, does not correlate with the modest changes in RNA Polymerase II (Pol II) occupancy and lack of changes in Pol II elongation observed in the two *Atxn7l3*^−/−^ cellular systems. These observations together indicate that deubiquitylation of histone H2Bub1 does not directly regulate global Pol II transcription elongation.

## Introduction

During mouse embryonic development, dynamic modifications of the chromatin are essential, as the loss of chromatin modifying enzymes, both writers and erasers, can lead to embryonic lethality, although with different severity ^1^. Histone H2B can be modified by the dynamic addition of a single ubiquitin (ub1) molecule on lysine 120 in mammals (H2Bub1). The deposition of mono-ubiquitin onto H2B is catalysed by the RNF20/RNF40 complex in mammals ^2–4^. The exact cellular function(s) of the H2Bub1 chromatin mark is not yet fully understood, however it was suggested that the H2Bub1 weakens DNA-histone interactions and therefore disrupts chromatin compaction ^5^. H2Bub1 mark was suggested to play a role in several DNA-related and epigenetically regulated processes, such as transcription, repair, replication, homologous recombination, as well as in mRNA processing and export ^6–15^. Indeed, chromatin immunoprecipitation coupled to sequencing studies revealed that H2Bub1 is found at gene bodies of expressed genes and is absent from non-expressed chromosomal regions, suggesting that H2Bub1 may be involved in transcription elongation ^16–20^. Intriguingly however, when H2Bub1 deposition was disrupted in mammalian cells by knock-down of RNF20 or knock-out of RNF40, the expression of only a small subset of genes was affected ^7, 12, 21^. H2Bub1 has also been implicated in histone crosstalk and shown to be a prerequisite for trimethylation of histone 3 lysine 4 (H3K4me3) around promoter regions both in yeast and mammalian cells ^22–27^.

H2Bub1 is erased by the deubiquitylation module (DUB) of the co-activator SAGA (Spt-Ada-Gcn5 acetyltransferase) complex ^28–31^, composed of the ubiquitin-specific protease 22 (USP22) and the ATXN7, ATXN7L3 and ENY2 adaptor proteins, needed for the full activity of USP22 enzyme ^32^. ATXN7L3 is critical for directing the DUB module substrate specificity towards H2Bub1 ^33^. In human cells, depletion of either ENY2 or ATXN7L3 adaptor protein resulted in a non-functional USP22 enzyme ^15, 19, 32, 34^. Two additional USP22-related ubiquitin hydrolases (called USP27X or USP51) can also interact with ATXN7L3 and ENY2, and deubiquitylate H2Bub1 independently of the SAGA complex ^34^. Thus, in mammalian cells the cellular abundance of H2Bub1 is regulated by the opposing activities of the ubiquitin E3 ligase complex, RNF20/RNF40, and three related DUB modules, each containing one of the homologous deubiquitylases: USP22, USP27X or USP51 ^34, 35^. These different DUB modules have also non-histone substrates ^36–43^. As *Usp22*^−/−^ mouse embryos die around embryonic day (E) E14.5 ^40, 44, 45^, the alternative USP27X- and/or USP51-containing DUB modules cannot completely fulfil the role of the USP22-containing DUB module. *Usp22*^−/−^ mouse studies suggest that USP22 is required to regulate apoptosis by deubiquitylating/stabilizing the SIRT1 histone deacetylase and by suppressing p53 functions during embryogenesis, and/or by regulating key signalling pathways crucial for mouse placenta vascularization ^40, 44^.

To better understand the role of USP22- and/or ATXN7L3-containing DUB modules *in vivo*, we have generated mice lacking USP22 or ATXN7L3. *Atxn7l3*^−/−^ embryos show developmental delay as early as E7.5, while *Usp22*^−/−^ embryos are normal at this stage, and die at E14.5 similarly to previous publications ^40, 44^. These results indicate that USP22 and ATXN7L3 are essential for normal development, however their requirements are not identical. H2Bub1 levels were only slightly affected in *Usp22*^−/−^ embryos, while in contrast they strongly increased in *Atxn7l3*^−/−^ embryos and derived cellular systems. The genome-wide increase of H2Bub1 retention in mESCs and MEFs lacking ATXN7L3 was investigated and the consequences of *Atxn7l3* loss of function on cellular homeostasis, differentiation, and RNA polymerase II (Pol II) transcription were analysed.

## Materials and Methods

### Generation and maintenance of *Usp22*^+/−^ and *Atxn7l3*^+/−^ mouse lines

*Usp22*^+/−^ and *Atxn7l3*^+/−^ mouse lines were generated at the Institut Clinique de la Souris (ICS, Illkirch, France) using mESCs containing the targeting constructs ordered from the International Knockout Mouse Consortium (IKMC), including the Knockout Mouse Programme (KOMP) repository (UC, Davis). In the *Usp22* targeting construct (*Usp22^tm1a(KOMP)Wtsi^*) a *LacZ* and *Neo* cassette were located in intron 1, flanked by *FRT* sequences, and *loxP* sequences were flanking exon 2 (Supplementary Fig. 1A). In the *Atxn7l3* targeting construct (*Atxn7l3^tm1.1(KOMP)Wtsi^*) a *LacZ* and *Neo* cassette were located in intron 2, flanked by *FRT* sequences, and the *LoxP* sequences were flanking exon 2 to exon 12 (Supplementary Fig. 1C). Chimeras were generated by injecting the C57BL/6 mESCs containing the targeting constructs into BALB/c blastocysts. Mice heterozygous for the targeting allele were crossed to a Cre-, or FLP-recombinase deleter strains, in order to generate the null alleles *Usp22*^−^ and *Atxn7l3*^−^, respectively. Then mice heterozygous for the null allele (*Usp22*^+/−^, or *Atxn7l3*^+/−^) were intercrossed to generate homozygous mutant embryos (*Usp22*^−/−^ or *Atxn7l3*^−/−^) as shown in Supplementary Fig. 1A and 1C. Investigators were not blinded for animal experimentation and no randomization was used as all the conditions were processed at the same time. Genotyping primers are shown in Supplementary Table 1, and examples of genotyping gels are shown in Supplementary Fig. 1B and 1D. *Atxn7l3*^+/−^ mice were maintained on a mixed B6D2 background. Animal experimentation was carried out according to animal welfare regulations and guidelines of the French Ministry of Agriculture and French Ministry of Higher Education, Research and Innovation.

### Generation and maintenance of *Atxn7l3*^−/−^ mESCs and *Atxn7l3*^−/−^ MEFs

To generate *Usp22*^−/−^, *Atxn7l3*^−/−^ and wild-type (WT) mESCs, timed matings between heterozygous mice were conducted, then at E3.5, pregnant females were sacrificed, uteri were flushed with M2 medium (Sigma-Aldrich), and individual blastocysts were transferred to wells of a 96-well plates pre-coated with 0.1% gelatin. Blastocysts were cultured and expanded in regular mESCs medium [DMEM (4.5 g/l glucose) with 2 mM Glutamax-I, 15% ESQ FBS (Gibco), penicillin, streptomycin, 0.1 mM non-essential amino acids, 0.1% ß-mercaptoethanol, 1500 U/mL LIF and two inhibitors (2i; 3 µM CHIR99021 and 1 µM PD0325901, Axon MedChem)]. After expansion, mESCs were genotyped and frozen.

To generate *Atxn7l3*^−/−^ and WT MEFs, timed matings between heterozygous mice were conducted, then at E10.5, pregnant females were sacrificed, and embryos were collected. The embryo yolk sacs were collected for genotyping, and the head and gastrointestinal tract were carefully dissected away from embryos. The remaining carcasses were transferred to individual 1.5 ml Eppendorf tubes, and 50 μl of 0.25% trypsin-EDTA (Gibco) was added and gently triturated 5 times to dissociate the embryos. The dissociated embryos were incubated in trypsin for 5 min at room temperature, then the trypsin was quenched with 500 μl of FCS. Cells were transferred to individual wells of a 6-well plate pre-coated with 0.1% gelatin and cultured in MEF medium (DMEM, 10% FCS, penicillin and streptomycin). Cells were visualized with an EVOS XL Core Cell Imaging System (#AMEX-1100, Thermo Fisher Scientific) using a LPlan PH2 10x / 0.25 objective.

### Protein extraction and Western blot assays

To extract histone proteins, embryos dissected at the indicated embryonic days, or about 5 x10^6^ cells (were lysed with 100 μl acidic extraction buffer (10 mM Hepes, pH 7.9, 1.5 mM MgCl_2_, 10 mM KCl, 0.5 mM DTT and 0.2 M HCl) freshly complemented with 1× Proteinase Inhibitor Cocktail (Roche) and 10 mM N-ethylmaleimide (Sigma-Aldrich). HCl was added to a final concentration of 0.2 M and incubated on an end-to-end rotator for 2 hours at 4°C. Following the incubation, cell extract was centrifuged at 20 800 x g for 10 min at 4°C, to pellet the acid insoluble material. A solution of 2 M Tris-HCl (pH 8.8) was added to neutralize the supernatant of the acidic extraction. Ten μl of the supernatant, containing histone proteins, were run on 4–12% gels (Bis-tris NuPAGE Novex, Life Technologies), then proteins were transferred and western blot assays were carried out by using standard methods. The following antibodies were used: anti-H3 (Abcam #ab1791) anti-H4 (Invitrogen 3HH4-4G8), anti-H2Bub1 (Cell Signaling Technology, #5546), anti-H3K4me3 (Abcam ab8580), anti-H3K9ac (Merck-Millipore #07-352), Peroxidase AffiniPure F(ab’)< Fragment Goat Anti-Mouse IgG, Fcγ fragment specific (Jackson ImmunoResearch #115-036-071) and Peroxidase AffiniPure Goat Anti-Rabbit IgG (H+L) (Jackson ImmunoResearch #115-035-144). Protein levels were quantified by ImageJ.

### Actin labelling

Cells were washed twice with 1x PBS, fixed with 4% PFA (Electron Microscopy Science) for 10 min at room temperature (RT). After fixation, cells were washed three times with 1x PBS, permeabilized with sterile 0.1% Triton X-100 in PBS for 20 min at RT, then washed three times in 1x PBS. Cells were incubated either with phalloidin conjugated to Alexa 488 dye (Phalloidin-iFluor 488, Abcam ab176753) following the manufacturer’s protocol, to label F-actin filaments, or with an anti-β-actin mouse monoclonal antibody (Sigma Aldrich, A5441) at a dilution of 1:1000 in 1x PBS with 10% FCS, overnight at 4°C. The following day, cells were washed three times with 1x PBS, then β-actin labelled cells were further incubated with secondary Alexa Fluor 488 goat anti-mouse Ig (Invitrogen #A-1101) at a dilution of 1:2000 in 1x PBS with 10% FCS for 1 hr at RT. The cells were washed three times with 1x PBS, then incubated in 20 mM Hoechst 3342 (Thermo Scientific) for 10 min at RT, before being washed three times with 1x PBS, then cells were covered with a coverslip coated in ProLong Gold mounting medium (Invitrogen). Pictures were taken using a Leica DM 4000 B upright microscope equipped with a Photometrics CoolSnap CF Color camera with a HCX PL S-APO 20x/0.50 objective.

### Colony formation assay and alkaline phosphatase staining

Four thousand mESCs were seeded on gelatin-coated 6-well plates in regular mESC medium (see above) to form colonies at low density. The medium was exchanged every two days for 6 days. Alkaline phosphatase (AP) activity test was performed using Red Substrate Kit, Alkaline Phosphatase (Vector Laboratories) according to the manufacturer’s instructions: mESC clones were washed with 1x cold PBS and fixed with 4% PFA for 10 min at RT. After fixation, cells were washed twice with H_2_O and incubated in 1 ml AP detection system for 30 min at RT in the dark. Then cells were washed twice with cold 1x PBS, and visualized with an EVOS XL Core Cell Imaging System (#AMEX-1100, Thermo Fisher Scientific) using a LPlan PH2 4x / 0.13 objective.

### Cell proliferation analysis

To determine cell proliferation, a total of 1×10^5^ mESCs per 6-well plate were seeded in regular mESC medium and 3×10^4^ passage 2-MEFs per 24-well plate were seeded in MEF medium. The medium was exchanged every two days. Cell numbers were counted with Countess cell counting chambers (Invitrogen). Statistical analyses were determined by a Mann-Whitney test (ns *p*>0.05; * *p* ≤ 0.05; ** *p* ≤ 0.01; *** *p* ≤ 0.001).

### Cell cycle analysis

Hundred thousand mESCs were fixed in 70% EtOH overnight at −4°C. After fixation, cells were treated with RNase A (100 μg/ml) (Thermo Fisher Scientific, #EN0531) and stained with propidium iodide (40 μg/ml) (Sigma Aldrich, #P-4170) for 30 min at 37°C. The acquisition of the DNA content was analysed on FACS CALIBUR (BD Sciences) flow cytometer. Quantitative results were analyzed by FlowJo software (BD Sciences).

### Apoptosis analysis using annexin-V staining

At the indicated incubation time, floating cells were collected in culture supernatants and adherent cells were harvested by trypsinization. After collection, cells were washed twice with cold 1X PBS, and about 2×10^5^ cells were resuspended in 100μl binding buffer (FITC Annexin V Apoptosis Detection Kit, Biolegend). Subsequently, 5μl FITC Annexin V (FITC Annexin V Apoptosis Detection Kit, Biolegend) and 10 μl propidium iodide was added to the cell suspension. Cells were gently vortexed and incubated in the dark for 15 min at RT. Thereafter, another 400 μl Annexin V binding buffer was added to each tube. Cells were analysed using a FACS CALIBUR (BD Sciences) flow cytometer. Dot plots were generated using the FlowJo software.

### Statistics

Statistical analysis of WT versus mutant samples comparison (cell proliferation and H2Bub1 density) was performed using non-parametric two-sided Wilcoxon rank sum test with continuity correction using R (version 5.3). No multiple comparisons were performed therefore no multiple correction were applied. Statistical results were expressed as *P* value or *P* value ranges (ns, *p* > 0.05; *, *p* ≤ 0.05; **, *p* ≤ 0.01; ***, *p* ≤ 0.001). Bar plot graphical data were represented as mean +/− SD and individual data points. Proliferation graphical data were represented as mean +/− SD. H2Bub1 density data were represented as box plots

Statistical analysis of the RNA-seq datasets was performed using a Wald test (DESeq2, see below).

### Enrichment of ubiquitylated peptides and quantification of H2Bub1 peptides

Forty million MEFs were harvested, washed three times with PBS and lysed in 1% SDS 0.1 M Tris pH 8 DTT 50 mM, 1× Proteinase Inhibitor Cocktail (Roche) and 50 mM PR-619 DUB inhibitor (UBPBio). The whole-cell lysate was precipitated with TCA and the pellet was washed twice with cold acetone then dried with Speed Vacuum system and weighted. Ten mg of the protein pellet was dissolved in 8 M urea, 5 mM TCEP then alkylated with 10 mM iodoacetamide. Sample was first digested with endoproteinase Lys-C (Wako) at a 1/500 enzyme/protein ratio (w/w) for 4h, then diluted four times before overnight trypsin digestion (Promega) at a 1:100 ratio. The resulting peptide mixture was desalted on C18 spin-column, quantified with Quantitative Colorimetric Peptide Assay (Thermo Fischer Scientific) and dried on Speed-Vacuum before enrichment with Ubiquitin Remnant Motif (K-ε-GG) Kit (Cell Signaling). After Sep Pak desalting, peptides were analyzed in triplicate using an Ultimate 3000 nano coupled in line with an Orbitrap ELITE (Thermo Scientific, San Jose California). Briefly, peptides were separated on a C18 nano-column with a 90 min linear gradient of acetonitrile and analyzed with Top 20 CID (Collision-induced dissociation) data-dependent acquisition method. Data were processed by database searching against Mus musculus Uniprot Proteome database (www.uniprot.org) using Proteome Discoverer 2.2 software (Thermo Fisher Scientific). Precursor and fragment mass tolerance were set at 7 ppm and 0.6 Da respectively. Trypsin was set as enzyme, and up to 2 missed cleavages were allowed. Oxidation (M, +15.995), GG (K, +114.043) were set as variable modification and Carbamidomethylation (C) as fixed modification. Proteins and peptides were filtered with False Discovery Rate <1% (high confidence). Lastly quantitative values were obtained from Extracted Ion Chromatogram (XIC) and exported in Perseus for statistical analysis ^46^.

### RNA-seq and ChIP-seq analyses

For RNA-seq, total RNA was extracted from mESCs and MEFs (3 biological replicates of WT and *Atxn7l3*^−/−^ for each cell types) using the NucleoSpin RNA isolation kit (Macherey-Nagel), according to manufacturer’s instructions. Libraries were generated from the purified RNA using TruSeq Stranded mRNA (Illumina) protocol. After checking the quality of the libraries with the Bioanalyser (Agilent), libraries were sequenced on the Illumina HiSeq 4000 at the GenomEast sequencing platform of IGBMC. The raw sequencing data generated reads were preprocessed in order to remove adapter, poly(A) and low-quality sequences (Phred quality score below 20), then were mapped to the mouse mm10 genome using STAR ^47^. After PCA analysis, one WT ESCs dataset was excluded because it did not cluster with the other biological replicates. Differentially expressed genes were measured using the DESeq2 package^48^. For the downstream analysis, only the transcripts which base median over 10 normalized reads (DESeq2 normalized reads divided by the median of the transcript length in kb) were considered. Using these criteria 16269 transcripts were expressed in mESCs, and 15084 transcripts were expressed in MEFs.

ChIP-seq experiments were performed using the protocol described in ^49^, with some minor modifications, including the use of 10 mM N-ethylmaleimide (NEM, Sigma-Aldrich) in all buffers and the use of either the anti-H2Bub1 antibody (MediMabs, NRO3), the anti-RPB1 CTD Pol II antibody (1PB 7G5 ^50^) from control and *Atxn7l3*^−/−^ mESCs and MEFs (n=1, each) or the anti-Ser2P CTD Pol II antibody (3E10 ^51^) from control and *Atxn7l3*^−/−^ mESCs (n=1, each). Briefly, mESCs or MEFs were fixed in 1% PFA for 10 min at RT, then the PFA was quenched with glycine at a final concentration of 125 mM for 5 min at RT. Cells were washed two times in 1× cold PBS, scraped, and pelleted. Nuclei were isolated by incubating cells with nuclear isolation buffer [50 mM Tris–HCl pH 8.0, 2 mM EDTA pH 8.0, 0.5% Nonidet P-40, 10% glycerol, Proteinase Inhibitor Cocktail (Roche), 10 mM NEM (and 1× PhosSTOP only in Pol II Ser2P ChIP samples)] for 10 min at 4°C with gentle agitation, followed by centrifugation at maximum speed to pellet the nuclei. Nuclei were resuspended in sonication buffer [0.1% SDS, 10 mM EDTA, 50 mM Tris-HCl pH 8.0, 1× Proteinase Inhibitor Cocktail, 10 mM NEM (and 1× PhosSTOP only in Pol II Ser2P ChIP samples)]. Then chromatin was sheared with the E220 sonicator (Covaris) and chromatin concentration was measured with the Qubit 3.0 (Thermo Fischer Scientific). Approximately of 50 µg of chromatin was used for Pol II or H2Bub1 ChIP, and 240 µg of chromatin was used for Pol II Ser2P ChIP which were diluted in ChIP dilution buffer [0.5% Nonidet P-40, 16.7 mM Tris–HCl pH 8.0, 1.2 mM EDTA, 167 mM NaCl, 1× Proteinase Inhibitor Cocktail, 10 mM NEM (and 1× PhosSTOP in Pol II Ser2P ChIP samples)]. Antibodies were incubated with the chromatin overnight with gentle agitation at 4°C. The next day, Dynabeads protein G magnetic beads (Invitrogen) were added for 1 hour, then were isolated and washed for 5 min at 4°C, three times with low salt wash buffer (0.1% SDS, 0.5% Nonidet P-40, 2 mM EDTA, 150 mM NaCl, 20 mM and Tris-HCl pH 8.0), three times with high salt wash buffer (0.1% SDS, 0.5% Nonidet P-40, 2 mM EDTA, 500 mM NaCl, 20 mM and Tris–HCl pH 8.0), and once with LiCl wash buffer (0.2 M LiCl, 0.5% Nonidet P-40, 0.5% sodium deoxycholate, 1 mM EDTA, 10 mM Tris-HCl pH8.0), then washed three times with TE buffer, then the beads were incubated in elution buffer (1% SDS, 0.1 M NaHCO_3_) at 65°C with shaking to elute complexes. Crosslinks were reversed with by adding NaCl at a final concentration of 0.2 M as well as 50 μg/ml RNase A at 65°C overnight and the following day the samples were treated with 20 µg Proteinase K, 26.6 µl of 1 M Tris–HCl pH 7.9, and 13.3 µl of 0.5 M EDTA incubated at 45°C for 1hr, and DNA was phenol/chloroform purified and precipitated. The precipitated DNA was used to generate libraries with the MicroPlex Library Preparation kit v2 (Diagenode) for ChIP-seq according to the manufacturer’s instructions. The samples were then sequenced on HiSeq 4000 with read lengths of 50 bp single end, reads were mapped to the mouse mm10 genome by the software Bowtie1. Samples were normalized (see below) and peak calling was performed using the MACS2 software.

### Bioinformatics tools and data-analysis methods

#### Normalization between ChIP-seq datasets

For H2Bub1 ChIP-seq samples to correct for the bias introduced by the differences in the sequencing depth among samples, the total reads present at 3115 intergenic regions far away from genes and larger than 100 kb were selected as described previously ^19^. These intergenic reads were used for the normalization of H2Bub1 ChIP-seq samples. We calculated the size factor of these intergenic regions for each sample using DESeq2 (version 1.16) ^52^. These size factors were used to normalize the H2Bub1 ChIP-seq data.

For Pol II ChIP-seq and Pol II Ser2P ChIP-seq samples, the total number of mapped reads were used for normalization. Twenty million of total reads were used to generate the .bed files for the seqMINER software analysis. The .bigwig files for the Integrative Genomics Viewer (IGV) visualization were generated with makeUCSCfile program in Homer package, the total number of reads was normalized to 10 million and the fragment length was normalized to 100 bp.

#### Calculation of density values

Density values were defined as follows: density = [(number of aligned reads in a region of interest) / (length of the region of interest in bp)] / (size factor x 10^−8^). For H2Bub1 datasets, we considered only the gene bodies of expressed genes containing at least 1 ChIP-seq read. Out of 16269 expressed genes in mESCs, 15467 contain at least 1 ChIP-seq read. Out of 15084 expressed genes in MEF cells, 14500 contain at least 1 ChIP-seq read (Supplementary Table 4).

#### Generation of average profiles and heat maps

Average profiles and k-means clustering were generated with the seqMINER program ^53^. The end of each aligned read was extended to 200 bp in the direction of the read. For the analyses around promoters, the tag density was extracted in a 2 kb window centred on each TSS. For average gene profiles, each gene body was divided into 160 equal bins (the absolute size depending on the gene length), 5 kb upstream and downstream were added. Moreover, 20 equally sized bins (250 bp/bin) were created upstream and downstream of genes. Densities were collected for each dataset in each bin.

#### Calculation of Pol II traveling ratio

Pol II pausing was based on the “Pausing Index” which is also referred to as “Traveling Ratio” ^54, 82^. We estimated the “Traveling Ratio” as the ratio of normalized Pol II ChIP-seq reads within the TSS region (–100 to +300 bps around TSS) to that in the gene body (TSS + 300 bps to TSS +2 kb), for genes expressed more than 100 normalized reads to the median size of transcript in kb and > 1 kb in length. Maximal distance and Kolmogorov & Smirnov test were analysed using R (version 3.5).

#### Code availability

Data and figures were generated using R (version 3.5). All custom code is available upon request.

### Data availability

All the datasets generated during the current study are available together in Gene Expression Omnibus (GEO) database under the accession number GSE153587. Individual RNA-seq data can be accessed at GSE153578 and ChIP-seq data at GSE153584.

## Results

### *In vivo* loss of ATXN7L3 results in a more severe phenotype than the loss of USP22

To compare the deubiquitylation requirement for USP22 and ATXN7L3 *in vivo*, we generated *Usp22*^+/−^ and *Atxn7l3*^+/−^ mice (Supplementary Fig. 1A-D). As heterozygous *Usp22^+/−^, or Atxn7l3*^+/−^ mice were phenotypically indistinguishable from their wild type littermates (Table 1 and 2), *Usp22^+/−^, or Atxn7l3*^+/−^ mice were intercrossed to obtain *Atxn7l3*^−/−^ and *Usp22*^−/−^ homozygous mutants. *Usp22*^−/−^ embryos started to resorb at E13.5 (Fig. 1Ah) and could not be observed after E14.5, similarly to what has been previously published ^40, 44, 45^ (Table 1, Fig. 1A). Similarly, no *Atxn7l3*^−/−^ pups could be retrieved at weaning (Table 2), however, analysis of *Atxn7l3*^+/−^ intercross litters collected at different stages of development revealed a more severe phenotype than *Usp22*^−/−^ mutants. A growth delay was already observed as early as E7.5 in *Atxn7l3*^−/−^ embryos (Fig. 1B), which did not turn at E9.5 (Fig. 1Bf). From E10.5 onwards, two classes of phenotype were observed: a severe and a mild, corresponding to 2/3 and 1/3 of the *Atxn7l3*^−/−^ embryos, respectively. No *Atxn7l3*^−/−^ embryos could be retrieved after E11.5 (Table 2). The mild class embryos were growth delayed (Fig. 1Bi and 1Bl) and in some instances blood pooling could be observed (Fig. 1Bh-Bl). The severe class embryos were smaller, failed to turn and displayed shortened trunk, abnormal head development, blood in the heart and enlarged pericardium (Fig. 1Bh and 1Bk). Our *in vivo* data demonstrate that loss of the DUB adaptor protein ATXN7L3 has a more severe effect on embryonic development than the USP22 enzyme, in agreement with published *in vitro* data ^34^.

**Fig. 1:**
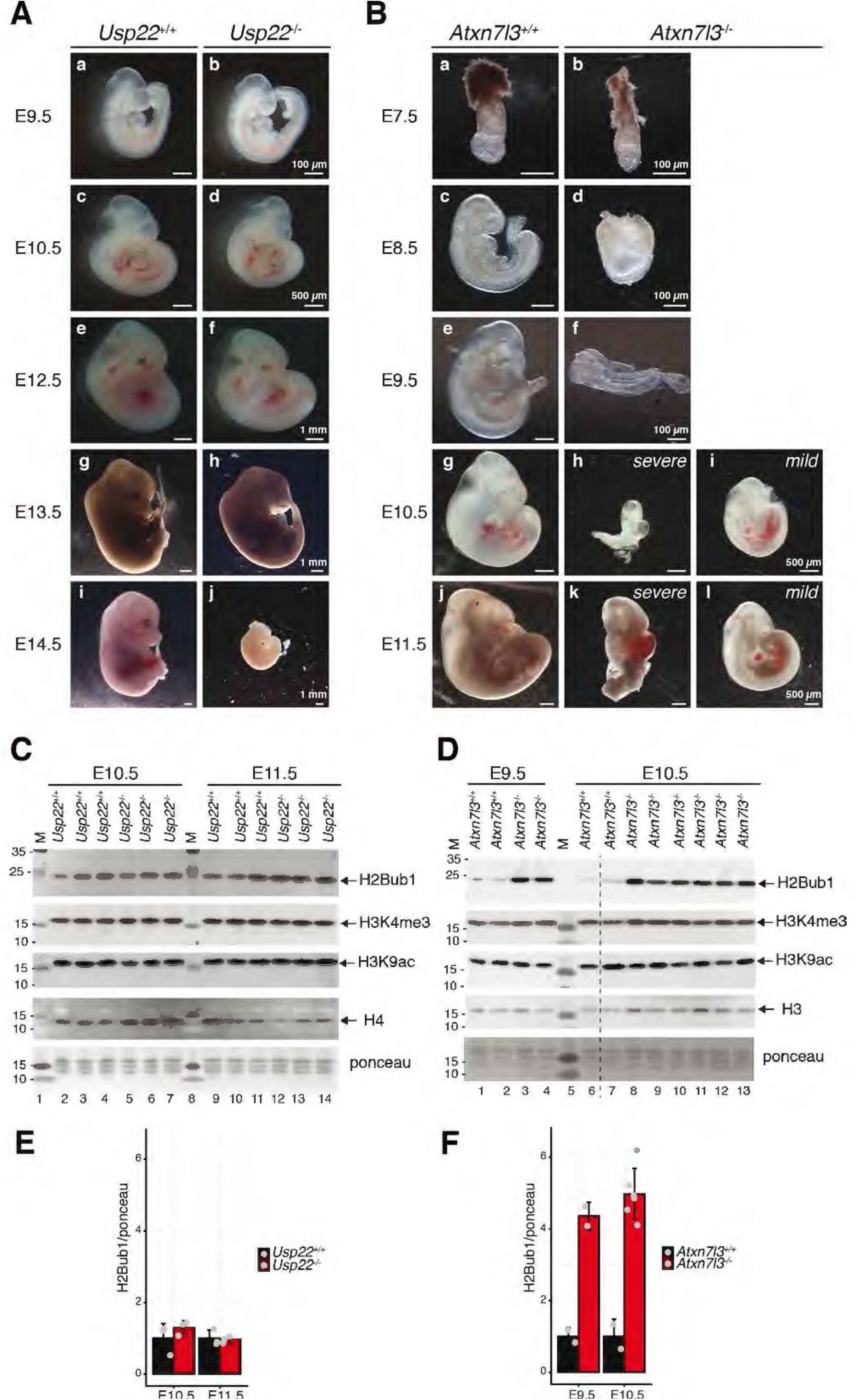
Loss of the SAGA DUB adaptor ATXN7L3 results in a more severe phenotype than loss of the DUB enzyme USP22. **A.** Comparison of *Usp22*^+/+^ and *Usp22*^−/−^ littermates from E9.5 to E14.5. **B.** Comparison of *Atxn7l3*^+/+^ and *Atxn7l3*^−/−^ littermates from E7.5 to E11.5. From E10.5 onwards, the *Atxn7l3*^−/−^ embryos can be categorized in 2 phenotypic classes; severe (h, k) and mild (i, l). **C-D.** Western blot analyses of E10.5 and E11.5 *Usp22*^+/+^ and *Usp22*^−/−^ (C), as well as E9.5 and E10.5 *Atxn7l3*^+/+^ and *Atxn7l3*^−/−^ (D) whole embryo lysates using anti-H2Bub1, anti-H3K4me3 and anti-H4 (C) or anti-H2Bub1, anti-H3K4me3 and anti-H3 (D) antibodies. A Ponceau staining view is displayed at the bottom of each panel. M: molecular weight marker (in kDa). The dotted line in (D) indicates where the blot was cut. Each lane represents a biological replicate. **E-F**. Western blot analyses shown in (C-D) were scanned and analysed densitometrically with ImageJ and the Ponceau normalized results are represented for each genotype.

**Table 1:** Offsprings from *Usp22*^+/−^ intercrosses

| Stage | *Usp22*<sup>+/+</sup> | *Usp22*<sup>+/-</sup> | *Usp22*<sup>-/-</sup> | Total | Number of litters |
| --- | --- | --- | --- | --- | --- |
| E9.5 | 3 (16.7%) | 9 (50%) | 6 (33.3%) | 18 | 2 |
| E10.5 | 5 (23.8%) | 11 (52.4%) | 5 (23.8%) | 21 | 3 |
| E12.5 | 8 (19.5%) | 21 (51.2%) | 12 (29.3%) | 41 | 5 |
| E13.5 | 4 (28.6%) | 7 (50%) | 3 (21.4%) | 14 | 2 |
| E14.5 | 6 (27.3%) | 10 (45.4%) | 6* (27.3%) | 22 | 3 |
| weaning | 93 (37.6%) | 154 (62.4%) | 0 (0%) | 247 | 37 |

\* dead embryo (no beating heart)

**Table 2:** Offsprings from *Atxn7l3*^+/−^ intercrosses

| Stage | *Atxn7l3*<sup>+/+</sup> | *Atxn7l3*<sup>+/-</sup> | *Atxn7l3*<sup>-/-</sup> | Total | Number of litters |
| --- | --- | --- | --- | --- | --- |
| E7.5 | 10 (47.6%) | 5 (23.8%) | 6 (28.6%) | 21 | 2 |
| E8.5 | 20 (31.2%) | 35 (54.7%) | 9 (14.1%) | 64 | 7 |
| E9.5 | 13 (25.5%) | 26 (51%) | 12 (23.5%) | 51 | 6 |
| E10.5 | 53 (28.8%) | 83 (45.1%) | 48 (26.1%) | 184 | 21 |
| E11.5 | 7 (28%) | 12 (48%) | 6 (24%) | 25 | 3 |
| E12.5 | 9 (47.4%) | 10 (52.6%) | 0 (0%) | 19 | 3 |
| weaning | 138 (44.7%) | 171 (55.3%) | 0 (0%) | 309 | 47 |

### *Atxn7l3*^−/−^ embryos show strong increase in global H2Bub1 levels

To investigate the importance of USP22 and ATXN7L3 on H2Bub1 deubiquitylation *in vivo*, we analysed global H2Bub1 levels from WT and *Usp22*^−/−^ embryos at E10.5, E11.5 and E12.5 stages (Fig. 1C, and data not shown), as well as from WT and *Atxn7l3*^−/−^ embryos at E9.5, or E10.5 stages (Fig. 1D). While minor changes (1.2 fold) were observed between controls and *Usp22*^−/−^ embryos lysates (Fig. 1C and 1E), a 4-5-fold increase in global H2Bub1 levels were observed in *Atxn7l3*^−/−^ embryo extracts (Fig. 1D and 1F), confirming similar observations made in human cancer cell lines ^32, 34^. Histone H3K4 trimethylation, and H3K9 acetylation were not affected in *Usp22*^−/−^, or *Atxn7l3*^−/−^ embryos (Fig. 1C and 1D). Thus, ATXN7L3 is required for the full activity of the three related DUB modules to regulate global H2Bub1 levels, whereas USP22-containing DUB module is less involved in genome-wide deubiquitylation of H2Bub1.

### *Atxn7l3*^−/−^ mESCs and *Atxn7l3*^−/−^ MEF-like cells show abnormal proliferation and phenotypes

As *Usp22*^−/−^ embryonic phenotypes have been already described ^40, 44^, we concentrated our analyses on *Atxn7l3*^−/−^ mutants. To determine the mechanistic outcome of perturbed DUB function(s), we derived mESCs and MEF-like cells from *Atxn7l3*^−/−^ embryos. As in the embryos, ATXN7L3 protein levels were undetectable in these *Atxn7l3*^−/−^ cellular systems (Supplementary Fig. 2A) and global H2Bub1 levels were significantly upregulated, by almost 4-5-fold in mESCs and about 7.5-8-fold in MEFs (Fig. 2A, 2B and Supplementary Fig. 2B).

**Fig. 2:**
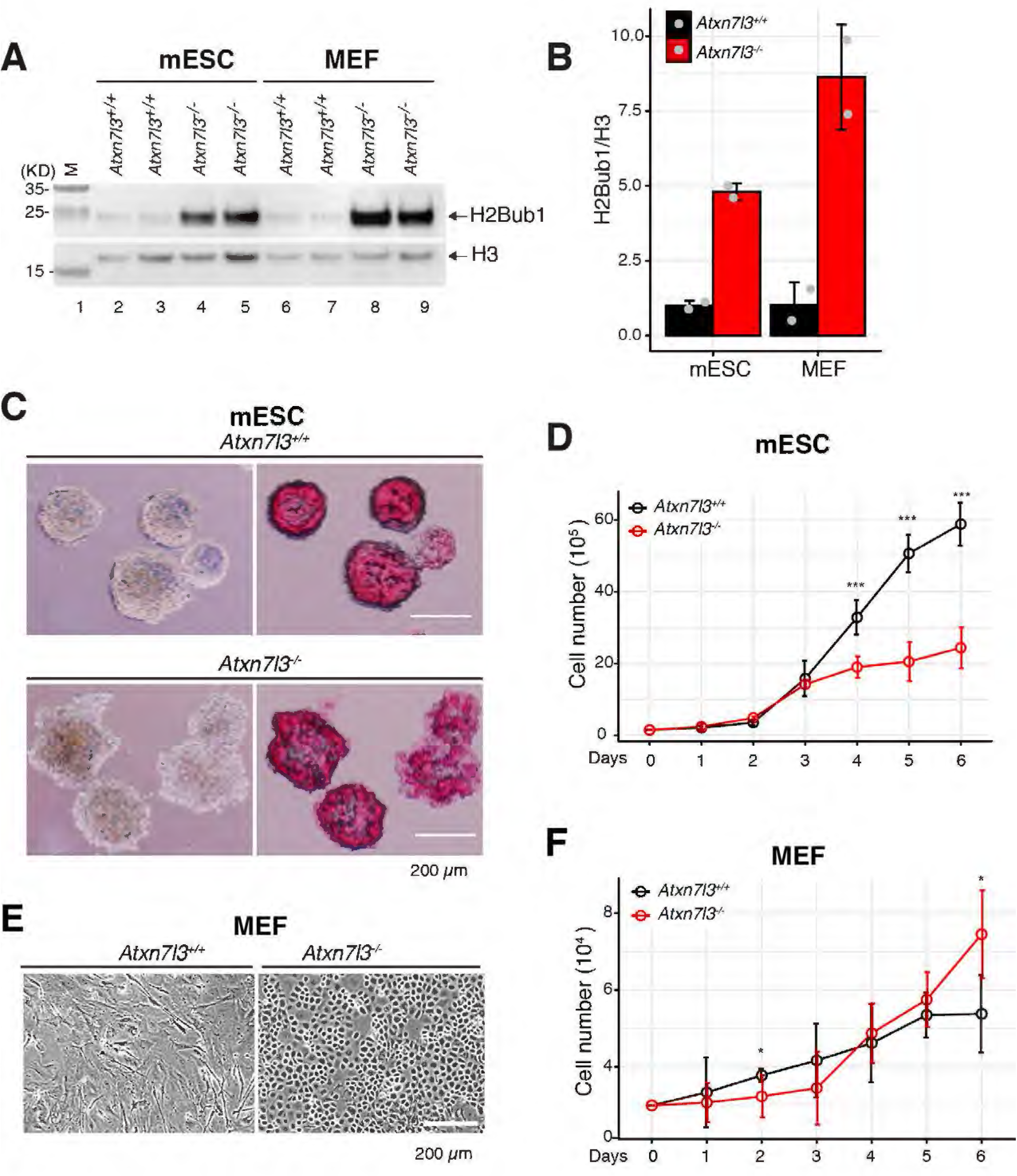
Primary *Atxn7l3*^−/−^ mESCs and *Atxn7l3*^−/−^ MEF-like cells show strong increase in H2Bub1 levels, abnormal proliferation and phenotypes. **A.** Western blot analysis of H2Bub1 levels in acidic histone extracts obtained from *Atxn7l3*^+/+^ or *Atxn7l3*^−/−^ mESC clones and *Atxn7l3*^+/+^ or *Atxn7l3*^−/−^ MEFs. Histone H3 western blot and ponceau stained membranes are shown as loading controls. Each lane represents a biological replicate. **B.** Quantification of H2Bub1 levels from (A) by using ImageJ. The y axis represents the fold change compared with WT cells. Histone H2Bub1 quantification was carried out with H3 normalization. Error bars indicate mean ±SD based on two biological replicates (represented by grey dots). **C.** *Atxn7l3*^+/+^ or *Atxn7l3*^−/−^ mESCs (3 biological replicates for each genotype) cultured in serum/LIF plus 2i medium for 6 days were either observed by phase contrast microscopy (left panels) or visualized by alkaline phosphatase staining (right panels). Scale bar, 200 μm. **D.** *Atxn7l3*^+/+^ or *Atxn7l3*^−/−^ mESCs cell proliferation was determined by cell counting at the indicated time points (2 biological and 3 technical replicates for each genotype). Error bars indicate mean ±SD based on two biological samples with three technical replicates for each. Statistical significance was calculated using two-sided Wilcoxon rank sum test with continuity correction (ns, *p* > 0.05; *, *p* ≤ 0.05; **, *p* ≤ 0.01; ***, *p* ≤ 0.001). **E.** Morphology of *Atxn7l3*^+/+^ and *Atxn7l3*^−/−^ MEFs derived from E10.5 embryos. Scale bar, 200 μm (> 12 biological replicates). **F.** MEF cell number was determined by cell counting at the indicated time points (2 biological and 3 technical replicates for each genotype). Error bars indicate mean ±SD based on two biological samples with three technical replicates for each. Statistical significance was calculated using two-sided Wilcoxon rank sum test with continuity correction (ns, *p* > 0.05; *, *p* ≤ 0.05; **, *p* ≤ 0.01; ***, *p* ≤ 0.001).

Alkaline phosphatase staining and expression of pluripotency markers, such as *Pou5f1, Sox2, Klf4, Nanog, Esrrb* and *Tfcp2l1* ^55^, were similar between *Atxn7l3*^−/−^ and control mESCs (Fig. 2C, and Supplementary Fig. 2C), indicating that the pluripotency of these cells was not significantly affected in absence of ATXN7L3. Similarly, when apoptotic cell death and non-synchronized cell cycle phase distribution were measured, no significant differences were detected when comparing WT and *Atxn7l3*^−/−^ mESCs (Supplementary Fig. 2D and 2E). However, we observed that *Atxn7l3*^−/−^ mESCs colonies were more irregular (Fig. 2C) and proliferated slower (Fig. 2D) compared to WT mESCs. Thus, ATXN7L3-regulated DUB activity may be necessary to facilitate efficient cell cycle progression and consequent cell proliferation, similarly to USP22 that is critical for progressing through G1 phase of the cell cycle ^37^.

In *Atxn7l3*^−/−^ MEFs, many cells had an abnormal round morphology (Fig. 2E, right panel) originating from clusters of cells that proliferated faster than elongated *Atxn7l3*^+/+^ MEFs (Fig. 2E). The round *Atxn7l3*^−/−^ cells were present in all MEFs generated from E10.5 *Atxn7l3*^−/−^ embryos (n>12 embryos). The proportion of round cells relative to elongated cells appeared to correlate with the severity of the phenotype. No significant differences were detected when comparing WT and *Atxn7l3*^−/−^ MEFs cell cycle phase distribution and apoptotic cell death (Supplementary Fig. 2F and 2G). However, *Atxn7l3*^−/−^ MEFs from passage 2 tended to proliferate somewhat slower for the first three days compared to WT MEFs, but then started to grow faster than WT MEFs (Fig. 2F).

Thus, ATXN7L3-linked DUB activity loss, and the resulting increased H2Bub1 levels do not result in severe phenotypic changes in *Atxn7l3*^−/−^ mESCs, but cause profound morphological changes and proliferation alterations in *Atxn7l3*^−/−^ MEF-like cells.

### *D*eregulation of gene expression is more severe in *Atxn7l3*^−/−^ MEFs than in *Atxn7l3*^−/−^ mESCs

To further characterize ATXN7L3-dependent DUB activity, we measured changes in steady state mRNA levels between *Atxn7l3*^+/+^ and *Atxn7l3*^−/−^ mESCs, as well as between *Atxn7l3*^+/+^ and *Atxn7l3*^−/−^ MEFs by carrying out RNA-seq analyses. We first verified whether the *Atxn7l3*^−/−^ MEF-like cells still belong to the MEF lineage in spite of their unusual morphology by comparing our MEFs RNA-seq results with 921 RNA-seq datasets from 272 distinct mouse cell types or tissues ^56^. This analysis indicated that the *Atxn7l3*^−/−^ MEF-like cells clustered together with *Atxn7l3*^+/+^ MEFs or fibroblasts (Supplementary Fig. 3C), suggesting that the *Atxn7l3*^−/−^ MEF-like cells belong to the fibroblast lineage.

Differential gene expression analysis between *Atxn7l3*^−/−^ and WT mESCs, or *Atxn7l3*^−/−^ and WT MEFs, showed that in both *Atxn7l3*^−/−^ samples there are significant numbers of genes which expression was up- or down-regulated (Fig. 3A and 3B, and Supplementary Fig. 4A and 4B). When compared to control cells, 1116 up-regulated and 810 down-regulated transcripts were identified in *Atxn7l3*^−/−^ mESCs, while 1185 up-regulated and 1555 down-regulated transcripts were found in the *Atxn7l3*^−/−^ MEFs (Fig. 3A and 3B). These observations suggest that out of approximately 15000 Pol II transcribed genes in mESCs (16269 transcripts), or in MEFs (15089 transcripts), ATXN7L3-linked DUB function regulates the transcription of only a subset of genes. In both cellular systems, down-regulated, up-regulated and unchanged gene sets were validated using RT-qPCR (Supplementary Fig. 2C, and Supplementary Fig. 3D and 3E). The fold change in differentially expressed gene was much more pronounced in *Atxn7l3*^−/−^ MEFs than in *Atxn7l3*^−/−^ mESCs (Fig. 3A and 3B), as in *Atxn7l3*^−/−^ MEFs about 151 transcripts changed their expression 32-fold or more (up and down), while in *Atxn7l3*^−/−^ mESCs only 2 genes changed their expression 32-fold (Fig. 3C). Moreover, when comparing the down- or up-regulated genes between *Atxn7l3*^−/−^ mESCs and *Atxn7l3*^−/−^ MEFs, only very few transcripts were similarly affected in the two cellular systems (Fig. 3D and 3E), suggesting that ATXN7L3-linked DUB activity regulates different subset of genes in the two cellular environments.

**Fig. 3:**
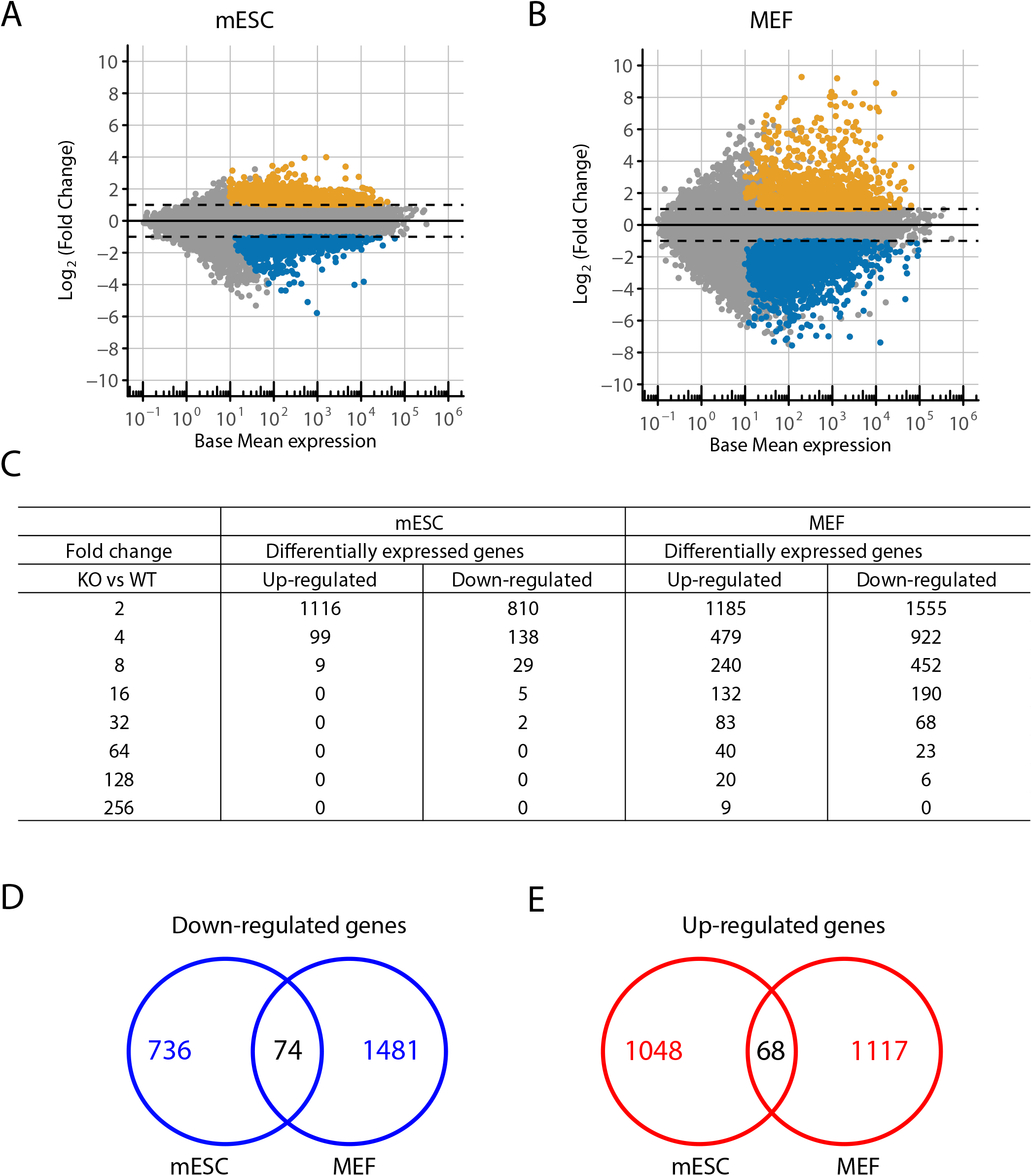
*Atxn7l3*^−/−^ mESCs and MEF-like cells show significant deregulation of transcription. **A-B.** MA-plots of RNA-seq data carried out on poly(A)^+^ RNA isolated from *Atxn7l3*^+/+^ and *Atxn7l3*^−/−^ mESCs (A, 2 and 3 biological replicates, respectively), or from *Atxn7l3*^+/+^ and *Atxn7l3*^−/−^ MEFs (B, 3 biological replicates, each). Log_2_ fold changes are shown versus Log_2_ mean expression signal. Differentially expressed genes were selected using the following thresholds: adjusted *p*-value ≤ 0.05, absolute value of Log_2_ fold change ≥ 1 and base median expression over 10 normalized reads to the median size of transcript in kb. Orange dots indicate up-regulated genes and blue dots indicates down-regulated genes. **C.** The number of significantly affected genes for *Atxn7l3*^−/−^ (KO)/*Atxn7l3*^+/+^ (WT) are represented for either mESCs or MEFs: adjusted *p*-value ≤ 0.5 and absolute value of fold change ≥ 2, 4, 8, 32, 64, 128, 256, separately. **D-E.** Venn diagrams indicate the overlap of down-regulated (**D**) and up-regulated (**E**) genes between mESCs and MEFs.

Gene ontology (GO) analyses of genes down-regulated in *Atxn7l3*^−/−^ mESCs revealed enrichment of GO categories linked to regulation of transcription, as well as cell differentiation, while in the up-regulated genes the GO categories “Metabolic processes”, and “Cell adhesion” were enriched (Supplementary Fig. 4C, 4D). Analyses of *Atxn7l3*^−/−^ MEFs indicated that genes involved in “Multicellular organism development”, and “Cell adhesion” were down-regulated, while genes belonging in “Metabolic” and “Immune system” processes were up-regulated (Supplementary Fig. 4E, 4F). Thus, ATXN7L3-related DUB activities regulate different subsets of genes in the two cellular systems.

### Cell adhesion and extracellular matrix genes are down-regulated in *Atxn7l3*^−/−^ MEFs

We further investigated the expression changes observed in the “Cell adhesion” GO category, since they could account for the unusual shape of the *Atxn7l3*^−/−^ MEFs. RNA-seq analyses indicated that a majority of genes coding for proteins belonging to this GO category: such as cadherins, catenins, collagens, and other cell adhesion molecules, were massively down-regulated in *Atxn7l3*^−/−^ MEFs compared to control MEFs (Fig. 4A). The deregulation of several of these genes was confirmed (Supplementary Fig. 3E).

**Fig. 4:**
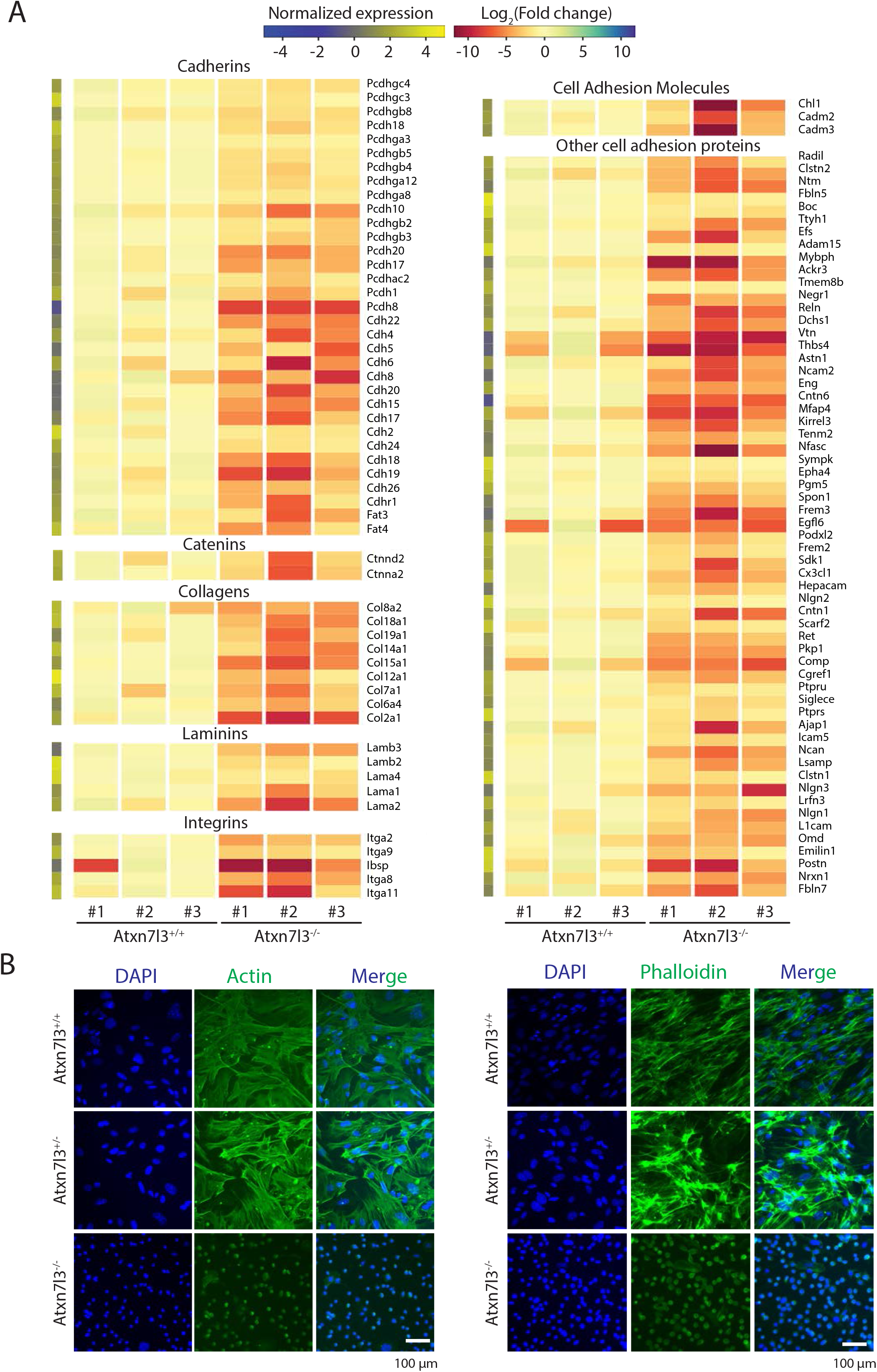
Cell adhesion genes are down-regulated in *Atxn7l3*^−/−^ MEFs. **A.** Heat map showing transcript levels belonging to the cell adhesion GO category from the three biological replicates of *Atxn7l3*^+/+^ and *Atxn7l3*^−/−^ MEFs for transcripts that are differentially expressed. Log_2_ of normalized expression is shown on the vertical column on the left. **B.** DAPI and immunofluorescence (IF) images of *Atxn7l3*^+/+^ and *Atxn7l3*^−/−^ MEFs stained with anti-β-actin antibody (left) and phalloidin (right) in MEF cells (n=1). The merge of DAPI and IF images is also shown. Scale bar: 100 μm.

We next analysed actin cytoskeletal proteins by fluorescence imaging. Using phalloidin staining, labelling F-actin filaments, and anti-β-actin immunofluorescence, we observed a massively reduced abundance of F-actin filaments and β-actin in *Atxn7l3*^−/−^ MEFs compared to WT MEFs (Fig. 4B), suggesting that loss of ATXN7L3 results in a down-regulation of cell adhesion complexes which in turn disrupt the actin cytoskeleton in MEFs.

### H2Bub1 levels increase in the gene bodies of *Atxn7l3*^−/−^ mESCs and *Atxn7l3*^−/−^ MEFs

To evaluate the changes in the genome-wide distribution of H2Bub1 in *Atxn7l3*^−/−^ mESCs, or *Atxn7l3*^−/−^ MEFs versus WT controls, chromatin immunoprecipitation coupled to high throughput sequencing (ChIP-seq) was performed using an anti-H2Bub1 antibody. The genomic distribution of H2Bub1 on several housekeeping genes was visualized using Integrative Genomics Viewer (IGV). H2Bub1 levels in both WT cell populations are relatively low, but highly increase in coding regions of both *Atxn7l3*^−/−^ mESCs and *Atxn7l3*^−/−^ MEFs, often showing a H2Bub1 enrichment peak downstream of the transcription start site (TSS) (Fig. 5A and 5B).

**Fig. 5:**
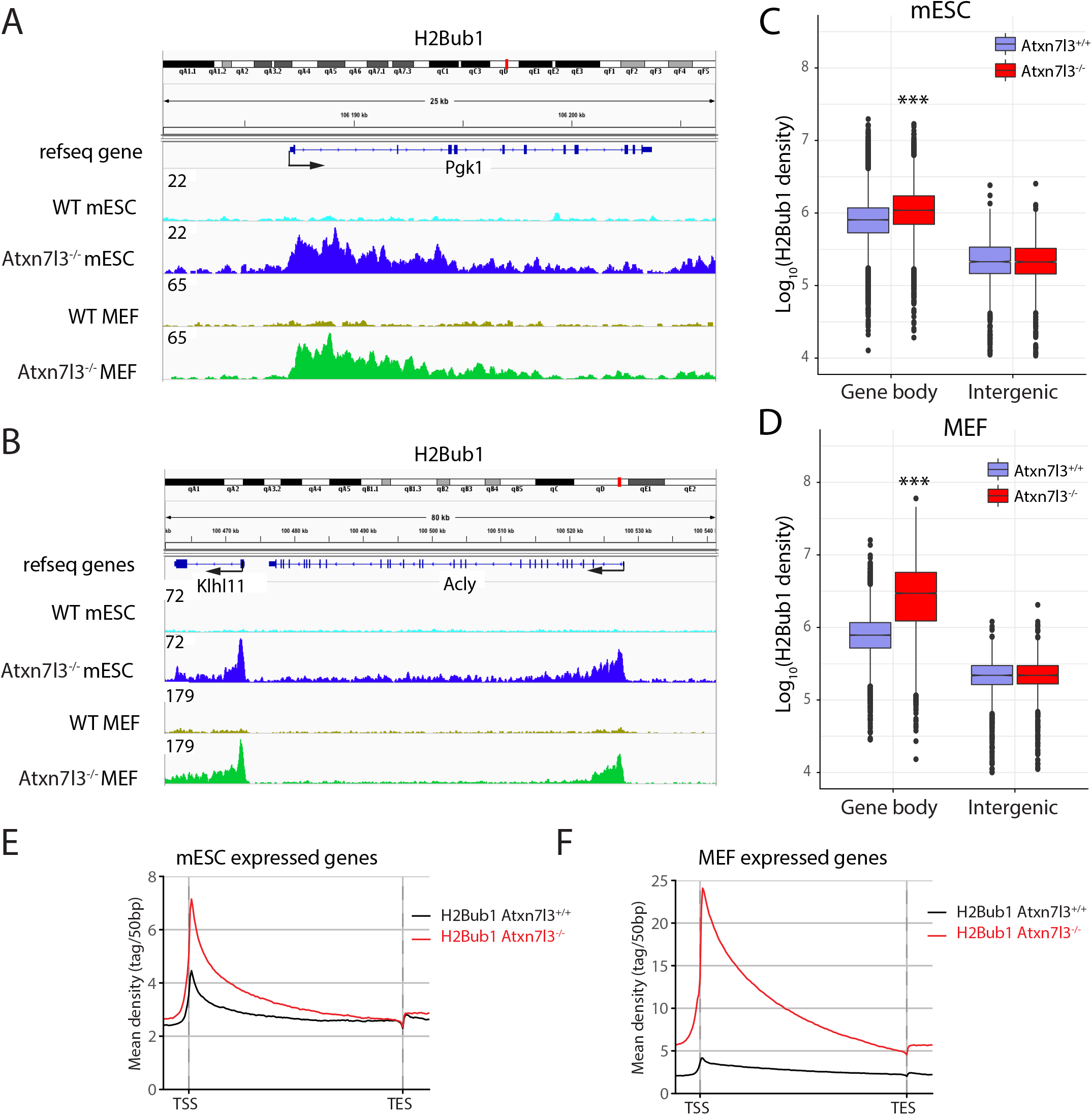
Histone H2Bub1 levels increase strongly in the gene bodies of both *Atxn7l3*^−/−^ mESCs and *Atxn7l3*^−/−^ MEFs. **A-B.** IGV genomic snapshots of H2Bub1 binding profiles (n=1) at three selected genes (*Pgk1*, *Klhl11* and *Acly*). Direction of the transcription is indicated by arrows. Group scaled tag densities on each gene either in mESCs, or in MEFs, are indicated on the left. **C-D.** Boxplots showing the Log_10_(H2Bub1 density) on the gene bodies of expressed transcripts or intergenic regions. Two-sided Wilcoxon rank sum test with continuity correction (***: *p*-value < 2.2e-16). **E-F.** Average metagene profiles showing H2Bub1 distribution on the bodies of expressed genes. 16269 expressed genes in mESCs (E) and 15084 expressed genes in MEFs cells (F) were chosen. TSS: transcription start site. TES: transcription end site. −5 kb region upstream of the TSS and +5 kb region downstream of the TES were also included in the average profile analyses.

To analyze quantitatively how the loss of the ATXN7L3-linked deubiquitylation activity changes H2Bub1 levels genome-wide, the presence of H2Bub1 over coding sequences of all annotated genes was normalized to intergenic regions and calculated. These analyses indicated that in *Atxn7l3*^−/−^ mESCs and in *Atxn7l3*^−/−^ MEFs the levels of H2Bub1 increase significantly over the gene body regions of either all expressed genes (Fig. 5C and 5D), or non-neighbouring expressed genes (after removing overlapping gene units within a 5 kb window 5’ and 3’; Supplementary Fig. 5A and 5B). In transcribed genes, we observed a 1.8-fold increase in H2Bub1 levels in *Atxn7l3*^−/−^ mESCs compared to WT controls, and a 6.5-fold increase in *Atxn7l3*^−/−^ MEFs (Fig. 5C and 5D, Supplementary Fig. 5C and 5D).

Next, metagene profiles of H2Bub1 spanning the entire transcribed regions and extending 5 kb upstream from TSSs and 5 kb downstream of the transcription end site (TES) were generated in *Atxn7l3*^−/−^ versus WT mESCs or MEFs (Fig. 5E and 5F). These profiles revealed a global increase of H2Bub1 over the whole transcribed region with an important enrichment in the region downstream from the TSS in *Atxn7l3*^−/−^ compared to WT mESCs (Fig. 5E). Similar results were obtained when we compared WT and *Atxn7l3*^−/−^ MEFs, however with a much stronger increase in H2Bub1 levels on the gene-body regions of *Atxn7l3*^−/−^ MEFs than in *Atxn7l3*^−/−^ mESCs (Fig. 5E and 5F). Moreover, our analyses on all expressed non-overlapping genes, up- or down-regulated in *Atxn7l3*^−/−^ cells (Supplementary Fig. 5E-5J) show that H2Bub1 levels increase strongly in all categories, except at genes down-regulated in *Atxn7l3*^−/−^mESCs (Supplementary Fig. 5F). These results show that ATXN7L3-linked DUB activity is responsible for the genome-wide deubiquitylation over the coding regions of expressed genes in mESCs and MEFs and suggest no general link between genome-wide H2Bub1 erasure defects and Pol II transcription.

### Pol II occupancy does not correlate with the H2Bub1 increase in *Atxn7l3*^−/−^ cells

To test whether the strong increase in H2Bub1 over the transcribed regions observed in the *Atxn7l3*^−/−^ cells would influence Pol II occupancy at promoters and/or in gene bodies, *Atxn7l3*^−/−^ mESCs and MEFs as well as control cells were subjected to ChIP-seq, using an antibody recognizing the non-modified C-terminal domain (CTD) of the largest subunit of Pol II (RPB1). Data obtained from *Atxn7l3*^−/−^ mESCs, and *Atxn7l3*^−/−^ MEFs at selected genes (Fig. 6A and 6B), or genome-wide (k-means clustering, Fig. 6C and 6D; and meta-gene plots, Fig. 6E and 6F) indicated that Pol II occupancy did not change dramatically, compared with the corresponding WT cells. Pol II occupancy at expressed genes was almost not affected at the TSS regions and slightly decreased in the gene body regions in both *Atxn7l3*^−/−^ cell types compared to WT cells (Fig. 6E-6F). In contrast, at most of Pol II occupied regions, H2Bub1 levels were highly increased in *Atxn7l3*^−/−^ mESCs and MEFs, compared to control cells (Fig. 5E and 5F). Next, we tested Pol II occupancy on genes, which were either down- or up-regulated by the loss of ATXN7L3-linked DUB activity (Fig. 3). Our metagene analyses on the up- and down-regulated gene categories in both *Atxn7l3*^−/−^ cell types suggest that Pol II occupancy changes tend to correlate with the observed changes in transcript levels, but not with changes in H2Bub1 levels (Supplementary Fig. 6A, 6B, 6E and 6F). As expected, we observed a complete loss of Pol II occupancy at highly down-regulated genes, or a strong increase in Pol II occupancy at highly up-regulated genes in *Atxn7l3*^−/−^ MEFs, compared to control cells (Supplementary Fig. 7). However, these totally opposite Pol II occupancy changes were often accompanied by a strong increase in H2Bub1 levels at these genes (Supplementary Fig. 7). These results together suggest that a strong global H2Bub1 increase in *Atxn7l3*^−/−^ cells do not globally deregulate RNA polymerase II occupancy at transcribed genes.

**Fig. 6:**
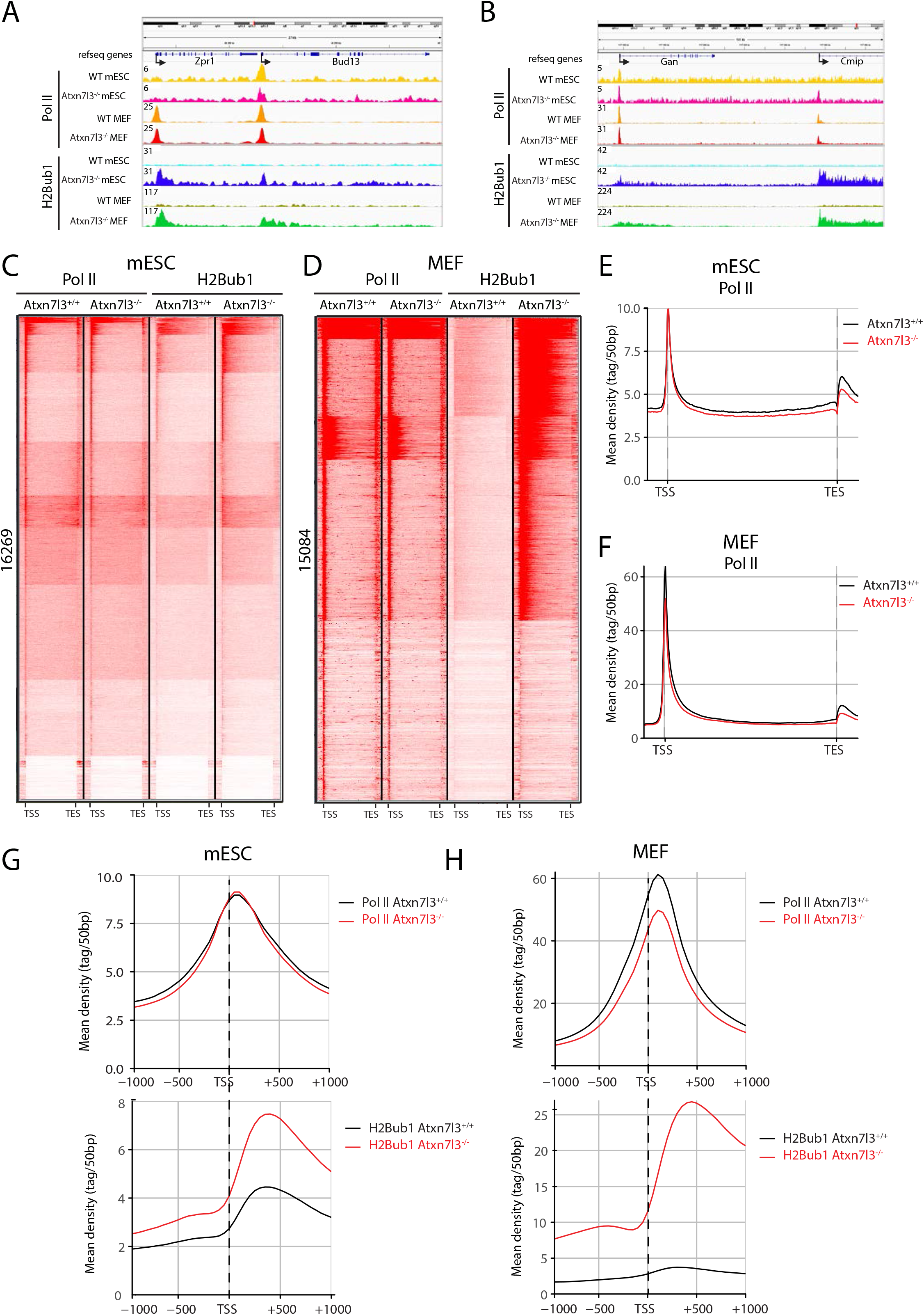
The modest genome-wide Pol II occupancy changes do not correlate with the strong H2Bub1 increases observed in the *Atxn7l3*^−/−^ mESCs or MEFs. **A-B.** IGV genomic snapshots of H2Bub1 and Pol II binding profiles (n=1) at four selected genes (*Zpr1*, *Bud13, Gan* and *Cmip*). Direction of the transcription is indicated by arrows. Group scaled tag densities on each gene either in mESCs, or in MEFs, are indicated on the left. **C-D.** K-means clustering showing the distribution of Pol II and H2Bub1 on genes expressed in mESCs (C, 16269 transcripts) and in MEFs (D, 15084 transcripts) (from −5 kb upstream from the TSS to + 5 kb downstream of the TES) in control and *Atxn7l3*^−/−^ mESC (C) and MEF (D). **E-F** Average metagene profiles showing Pol II distribution on bodies of expressed genes (from −5 kb upstream from the TSS to + 5 kb downstream of the TES) in control and *Atxn7l3*^−/−^ mESCs (E) and MEFs (F). **G-H.** Average profiles depicting Pol II and H2Bub1 distribution around the TSS (TSS −1 kb / +1 kb) of expressed genes in control and *Atxn7l3*^−/−^ mESCs (G) and MEFs (H).

### Paused Pol II and H2Bub1 peaks downstream of the TSSs do not overlap

Next, we analyzed whether promoter proximal Pol II peaks observed at transcribed genes around the +60 bp region ^57–59^, would overlap with the H2Bub1 peak observed downstream of the TSSs both in WT and *Atxn7l3*^−/−^ cells. As expected meta-gene analyses around the TSSs showed that in both mESCs and MEFs (WT and *Atxn7l3*^−/−^) Pol II peaks gave the highest signal at around the +60 region (Fig. 6G and 6H). In contrast, H2Bub1 density in WT and *Atxn7l3*^−/−^ mESCs and *Atxn7l3*^−/−^ MEFs is low in the +60 regions and reaches its maximum more downstream, in the +300 bp region (Fig. 6G and 6H). Importantly, Pol II accumulation at the pause site was not influenced by the large increase of H2Bub1 in *Atxn7l3*^−/−^ cells, suggesting that H2Bub1 deubiquitylation by the ATXN7L3-dependent DUB module(s) may not regulate promoter proximal pausing of Pol II, and/or Pol II turnover.

### Pol II elongation rates are not changed by H2Bub1 increase in *Atxn7l3*^−/−^ cells

As transcription elongation-linked Ser2 phosphorylation (Ser2P) of Pol II peaks downstream of the TSS and then increases again gradually in the gene body region ^60^, we performed anti-Pol II-Ser2P ChIP-seq in the *Atxn7l3*^−/−^ mESCs, compared to WT cells. Our genome-wide analyses of Pol II-Ser2P occupancy on expressed genes either by k-means clustering or by metagene profiling indicated no detectable changes around the TSS regions, but a slight (1.13-fold) increase of the Pol II-Ser2P signal towards the TES and downstream of it (Fig. 7A and 7B). K-means clustering analyses distinguished genes with a strong Pol II-Ser2P signal around their TESs in WT mESCs (cluster 1, Fig. 7A) and genes devoid of Pol II-Ser2P enrichment around their TESs (cluster 2, Fig. 7A). Metagene profiles of each cluster indicated that the increase of Pol II-Ser2P signal downstream of the TES in *Atxn7l3*^−/−^ mESCs was more pronounced at genes of cluster 1 than at cluster 2 (1.4-fold, Fig. 7A, 7C and 7D), suggesting potential defects in Pol II transcription, elongation rates and/or termination on genes belonging to cluster 1.

**Fig. 7:**
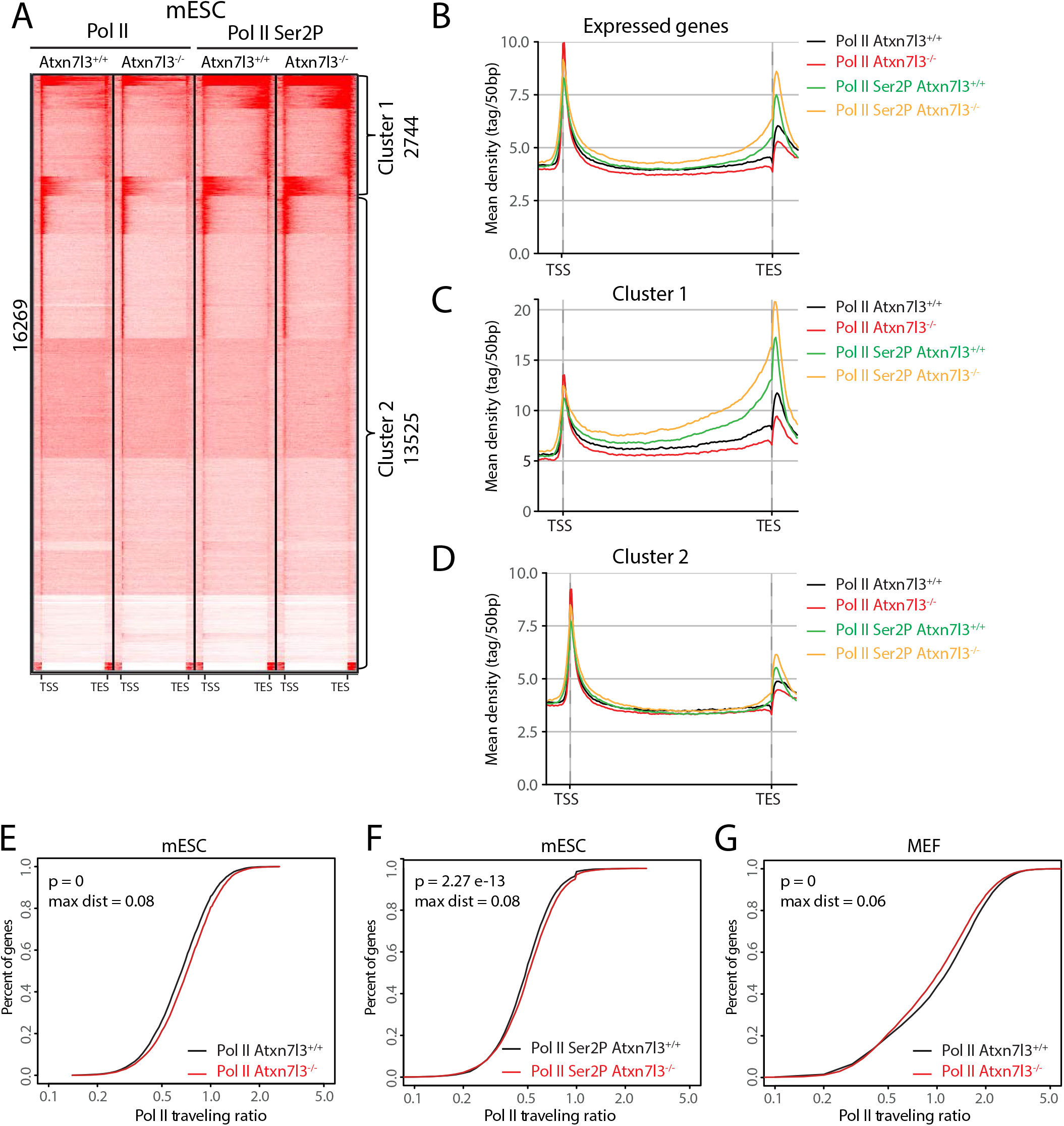
Pol II elongation rates are not changed by H2Bub1 increase in the gene-body regions of *Atxn7l3*^−/−^ cells. **A.** K-means clustering showing the distribution of Pol II and Pol II Ser2P (n=1) on 16269 expressed genes in mESCs (from −5 kb upstream from the TSS to + 5 kb downstream of the TES) in control and *Atxn7l3*^−/−^ mESC. Two clusters have been separated (as indicated) based on the Pol II Ser2P signal accumulation around the TES region in WT ESCs. **B-D** Average metagene profiles showing Pol II and Pol II Ser2P distribution on bodies of expressed genes (B), Cluster 1 (C) or Cluster 2 (D) (from −5 kb upstream from the TSS to + 5 kb downstream of the TES) in WT and *Atxn7l3*^−/−^ mESCs. The color code is indicated on the right of the panels. **E-G.** Pol II (E) and Pol II Ser2P (F) travelling ratios were calculated in WT and *Atxn7l3*^−/−^ mESC, and Pol II (G) traveling ratio was calculated in WT and *Atxn7l3*^−/−^ MEFs for genes expressed more than 100 normalized reads to the median size of transcript in kb. Kolmogorov & Smirnov test p values and maximal distance (max dist) are indicated.

Thus, we determined whether Pol II elongation rates in gene bodies was altered in *Atxn7l3*^−/−^ mESCs or MEFs, compared to WT cells. To this end we measured Pol II travelling ratios ^54, 61^. Pol II and Pol II-Ser2P travelling ratios in *Atxn7l3*^−/−^ mESCs, or Pol II traveling ratio in *Atxn7l3*^−/−^ MEFs, compared to WT cells, showed only very minor changes on expressed genes (Fig. 7E-7G), suggesting that Pol II elongation rates were not significantly changed in *Atxn7l3*^−/−^ mESCs or MEFs.

## Discussion

### Loss of the DUB adaptor ATXN7L3 results in a more severe phenotype than the loss of the DUB enzyme of SAGA, USP22

The relative abundance and function of the various DUB complexes, their redundant activities and/or compensatory mechanisms, in different cell types, at various stages of mouse embryogenesis has not been explored. Koutelou et al. (2019) ^44^ revealed that USP22 is essential for placental development. Consistent with our findings, *Usp22*^−/−^ embryos developed normally up to E12.5, but then die around E13.5-E14.5. It has been reported that *Usp22* is expressed ubiquitously in the embryo and hippomorphic *Usp22*^lacZ/lacZ^ mice have a reduced body size and weight ^45^. The absence of a strong morphological phenotype in the *Usp22*^−/−^ embryos before E13.5 suggests that many key early developmental processes do not require USP22, or that USP22 function can be compensated by other USPs, such as USP27X, USP51. It is however remarkable that placental development in *Usp22*^−/−^ embryos cannot be compensated by other USPs, suggesting a possible direct requirement of the SAGA complex in placental development.

On the other hand, no compensation is expected in *Atxn7l3*^−/−^ embryos as the absence of ATXN7L3 is supposed to inactivate all SAGA-related DUB complexes ^34^. Indeed, the *Atxn7l3*^−/−^ phenotype is more severe than that of *Usp22*^−/−^, occurring as early as E7.5. Although it is not known whether the H2Bub1 deubiquitylation is linked to *Usp22*^−/−^ or *Atxn7l3*^−/−^ embryos phenotypes, it is interesting to note that there is a parallel between the severity of the phenotypes and the changes in H2Bub1 levels. E10.5 *Usp22*^−/−^ embryos are normal and their genome-wide histone H2Bub1 levels do not increase (Fig. 1C and 1E), while in contrast E10.5 *Atxn7l3*^−/−^ embryos are seriously affected and their H2Bub1 levels increase 4-5-fold (Fig. 1D and 1F).

We observed two categories of *Atxn7l3*^−/−^ mutants: i) the severely affected embryos (2/3^rd^) which are growth retarded, fail to turn and display shortened trunk and abnormal head development; ii) the mildly affected embryos (1/3^rd^), which do turn and only display mild growth delay. It is conceivable that ATXN7L3 is involved in embryo patterning as for example, Nodal signalling mutant embryos, which are defective in early patterning of the primitive streak, also fail to turn ^62^. Nevertheless, the fact that some *Atxn7l3*^−/−^ embryos escape the severe phenotype suggest that ATXN7L3 and the corresponding DUB module(s) could be involved in a developmental checkpoint control at the time of embryo turning. Remarkably, all *Atxn7l3*^−/−^ embryos die around E11.5, and in addition to placental defects, they exhibit cardiovascular defects, as enlarged pericardium and blood pooling in the heart are observed in the severely affected *Atxn7l3*^−/−^ embryos. Thus, the comparison of the *Usp22*^−/−^ and *Atxn7l3*^−/−^ embryo phenotypes suggest that the defects observed in *Usp22*^−/−^ embryos could be compensated until E13.5 in the absence of USP22 by the activity of USP27X- and/or USP51-containing DUBs, which would require ATXN7L3 and ENY2 cofactors. Such compensation would not happen in *Atxn7l3*^−/−^ embryo, as in the absence of ATXN7L3 all three related DUBs would be inactive.

The comparisons of *Atxn7l3* regulated genes (RNA-seq from our study) with SAGA-bound genes (anti-TAF6L ChIP-seq; ^63^) in mESCs suggest that only a minority (< 8%) of SAGA-bound genes are regulated (up or down) by the loss of ATXN7L3-linked DUBs. This further suggests that the USP27x- and/or USP51-containing DUBs may have also SAGA-independent gene regulatory functions.

In conclusion, our results showing that *Usp22*^−/−^ embryo phenotypes are less severe agree with the biochemical findings suggesting that in *Usp22*^−/−^ cells the activity of only one of the three related DUB modules is eliminated. In contrast, in the *Atxn7l3*^−/−^ embryos the activities of all three related DUB modules are eliminated, thus, causing a more severe phenotype. It is also possible that ATXN7L3 loss may influence the epithelial-mesenchymal transition during gastrulation. The fact that *Atxn7l3*^−/−^ embryos survive until E11.5, suggests that none of these three related DUBs would play an essential role before this embryonic stage, and that also H2Bub1 deubiquitylation is not essential for Pol II transcription before this developmental stage.

### Histone H2Bub1 deubiquitylation is not linked to global RNA polymerase II transcription

Although H2B monoubiquitylation has been linked to increased transcription, transcription elongation, DNA replication, mitosis, and meiosis ^64^, how this histone modification and the erasing of this mark function is not well understood. In *Saccharomyces cerevisiae* H2B mutant, K123R (which cannot be ubiquitylated) the expression of only a low number of genes (about 300-500) is affected ^65^, suggesting that ubiquitylation and deubiquitylation may not have global transcriptional regulatory functions in yeast. Nevertheless, it has been suggested that H2Bub1 stimulates FACT-mediated displacement of a H2A/H2B dimer which in turn would facilitate the passage of Pol II through the nucleosome ^16^ and that H2Bub1 would be required for efficient reassembly of nucleosomes behind the elongating Pol II ^66, 67^. Contrary, it was reported that the effect of H2Bub1 on nucleosome stability is relatively modest ^68^.

If H2Bub1 deposition (by RNF20/RNF40), or H2Bub1 erasure (by the ATXN7L3-containing DUB modules) would carry out opposite genome-wide actions, their gene regulatory actions would result in mirroring effects. However, when comparing RNA-seq data from *Rnf40* knock-out MEFs ^12^ with our RNA-seq obtained from *Atxn7l3*^−/−^ MEFs, we did not observe any anti-correlation between the regulated genes in these two datasets (the Pearson correlation coefficient is −0.042, Supplementary Fig. 8), suggesting that the H2Bub1 deposition and erasure are not (only) carrying out opposite functions on the transcribed genome.

Contrary to H2B monoubiquitylation, it is much less well understood whether H2Bub1 deubiquitylation would be a process significantly impacting Pol II transcription. Previously, by using an *ATXN7L3* knock-down strategy in HeLa cells we showed that the ATXN7L3-related DUB activities are directed toward the transcribed region of almost all expressed genes, but correlated only poorly with gene expression ^19^. Our present results indicate that impairment of H2Bub1 deubiquitylation does not directly impact transcription initiation and/or elongation, because while we observe a massive H2Bub1 retention at almost every expressed gene in both *Atxn7l3*^−/−^ mESCs and *Atxn7l3*^−/−^ MEFs, Pol II and Pol II-Ser2P occupancy were only slightly impacted and only limited subsets of genes changed expression in both cellular systems (Fig. 3, 5-7). In addition, when comparing nascent and steady state mRNA changes of selected transcripts (Supplementary Figure 3 D) in WT ESCs and *Atxn7l3*^−/−^ ESCs, we did not observe any significant differences (data not shown). Moreover, in both cellular systems the lack of correlation between global H2Bub1 increase and consequent inhibition of global transcription suggests that H2Bub1 deubiquitylation does not directly regulate Pol II transcription. In agreement, the H3K4me3 chromatin mark present at the TSSs of active genes in eukaryotes, did not change in *Usp22*^−/−^ or in *Atxn7l3*^−/−^ embryos, in spite of the fact that in *Atxn7l3*^−/−^ embryos the H2Bub1 levels were increased by 4-5-fold (Fig. 1C and 1D). Similarly, global H3K9ac levels do not change in *Usp22*^−/−^ or in *Atxn7l3*^−/−^ embryos (Fig. 1C and 1D). Thus, our study corroborates other recent studies demonstrating catalytic-independent functions of chromatin modifying complexes in mouse ES cells ^69–71^.

Our results also suggest that the dynamic erasure of the H2Bub1 mark does not seem to influence global Pol II recruitment, pre-initiation complex formation at promoters, promoter proximal pausing or Pol II elongation rates (Fig. 6G, 6H and Fig. 7E-7G). However, Ser2 phosphorylation of Pol II in cluster 1 downstream of the TES regions in *Atxn7l3*^−/−^ ESCs was more pronounced, compared to WT cells (about 1.4-fold, Fig. 7A-7C), suggesting a possible compensatory «sensing» mechanism by the kinases regulating Pol II transcription elongation.

Whether the observed embryo and cellular phenotypes in the *Atxn7l3*^−/−^ embryos can be directly linked to increased H2Bub1 levels in specific transcribed regions, and/or to deubiquitylation failures of other ubiquitylated protein targets, will need to be further investigated in future.

## Acknowledgements

We thank all members of the Tora lab for protocols, thoughtful discussions and suggestions, especially V. Hisler for help with mice dissection, JC. Andrau for advice on Ser2P ChIP-seq, C. Hérouard and M. Jung from the GenomEast platform [France Génomique consortium (ANR-10-INBS-0009], for library preparation, sequencing and analyses; C. Ebel and M. Philipps for help with FACS, the IGBMC histology platform, the IGBMC cell culture facility and S. Falcone, M. Poirot and F. Memedov of the IGBMC animal facility for animal care taking. This study was supported by grants from European Research Council (ERC) (ERC-2013-Advanced grant 340551, Birtoaction), Agence Nationale de la Recherche (ANR) PICen-19-CE11-0003-02 and EpiCAST-19-CE12-0029-01 grants, NIH 1R01GM131626-01 grant (to LT) and ANR-18-CE12-0026 grant (to DD), by IGBMC International PhD program LABEX fellowship (to FW); and by the IdEx-University of Strasbourg PhD program and by the ‘Fondation pour la Recherche Médicale’ (FRM) association (FDT201904008368) (to VF), and an ANR-10-LABX-0030-INRT grant, under the frame program Investissements d’Avenir ANR-10-IDEX-0002-02.

## Notes

**Conflict of interest:** The authors declare no conflict of interest.

**Funding**: This study was supported by grants from European Research Council (ERC) (ERC-2013-Advanced grant 340551, Birtoaction), Agence Nationale de la Recherche (ANR) PICen-19-CE11-0003-02 and EpiCAST-19-CE12-0029-01 grants, NIH 1R01GM131626-01 grant (to LT) and ANR-18-CE12-0026 grant (to DD), fellowships by the IdEx-University of Strasbourg international PhD program and by the ‘Fondation pour la Recherche Médicale’ (FRM) association (FDT201904008368) (to VF) and an ANR-10-LABX-0030-INRT grant, a French State fund managed by the ANR under the frame program Investissements d’Avenir ANR-10-IDEX-0002-02.

https://www.ncbi.nlm.nih.gov/geo/query/acc.cgi?acc=GSE153587

### Competing Interest Statement

The authors have declared no competing interest.

## References

1. Festuccia N, Gonzalez I, Navarro P. The Epigenetic Paradox of Pluripotent ES Cells. J Mol Biol 2017, 429(10): 1476–1503.

2. Osley MA. Regulation of histone H2A and H2B ubiquitylation. Brief Funct Genomic Proteomic 2006, 5(3): 179–189.

3. Zhu B, Zheng Y, Pham AD, Mandal SS, Erdjument-Bromage H, Tempst P, et al. Monoubiquitination of human histone H2B: the factors involved and their roles in HOX gene regulation. Mol Cell 2005, 20(4): 601–611.

4. Shiloh Y, Shema E, Moyal L, Oren M. RNF20-RNF40: A ubiquitin-driven link between gene expression and the DNA damage response. FEBS Lett 2011, 585(18): 2795–2802.

5. Fierz B, Chatterjee C, McGinty RK, Bar-Dagan M, Raleigh DP, Muir TW. Histone H2B ubiquitylation disrupts local and higher-order chromatin compaction. Nat Chem Biol 2011, 7(2): 113–119.

6. Minsky N, Shema E, Field Y, Schuster M, Segal E, Oren M. Monoubiquitinated H2B is associated with the transcribed region of highly expressed genes in human cells. Nat Cell Biol 2008, 10(4): 483–488.

7. Shema E, Tirosh I, Aylon Y, Huang J, Ye C, Moskovits N, et al. The histone H2B-specific ubiquitin ligase RNF20/hBRE1 acts as a putative tumor suppressor through selective regulation of gene expression. Genes Dev 2008, 22(19): 2664–2676.

8. Trujillo KM, Osley MA. A role for H2B ubiquitylation in DNA replication. Mol Cell 2012, 48(5): 734–746.

9. Kari V, Shchebet A, Neumann H, Johnsen SA. The H2B ubiquitin ligase RNF40 cooperates with SUPT16H to induce dynamic changes in chromatin structure during DNA double-strand break repair. Cell Cycle 2011, 10(20): 3495–3504.

10. Moyal L, Lerenthal Y, Gana-Weisz M, Mass G, So S, Wang SY, et al. Requirement of ATM-dependent monoubiquitylation of histone H2B for timely repair of DNA double-strand breaks. Mol Cell 2011, 41(5): 529–542.

11. Nakamura K, Kato A, Kobayashi J, Yanagihara H, Sakamoto S, Oliveira DV, et al. Regulation of homologous recombination by RNF20-dependent H2B ubiquitination. Mol Cell 2011, 41(5): 515–528.

12. Xie W, Nagarajan S, Baumgart SJ, Kosinsky RL, Najafova Z, Kari V, et al. RNF40 regulates gene expression in an epigenetic context-dependent manner. Genome Biol 2017, 18(1): 32.

13. Vitaliano-Prunier A, Babour A, Herissant L, Apponi L, Margaritis T, Holstege FC, et al. H2B ubiquitylation controls the formation of export-competent mRNP. Mol Cell 2012, 45(1): 132–139.

14. Pirngruber J, Shchebet A, Schreiber L, Shema E, Minsky N, Chapman RD, et al. CDK9 directs H2B monoubiquitination and controls replication-dependent histone mRNA 3’-end processing. EMBO Rep 2009, 10(8): 894–900.

15. Evangelista FM, Maglott-Roth A, Stierle M, Brino L, Soutoglou E, Tora L. Transcription and mRNA export machineries SAGA and TREX-2 maintain monoubiquitinated H2B balance required for DNA repair. J Cell Biol 2018, 217(10): 3382–3397.

16. Pavri R, Zhu B, Li G, Trojer P, Mandal S, Shilatifard A, et al. Histone H2B monoubiquitination functions cooperatively with FACT to regulate elongation by RNA polymerase II. Cell 2006, 125(4): 703–717.

17. Chandrasekharan MB, Huang F, Sun ZW. Histone H2B ubiquitination and beyond: Regulation of nucleosome stability, chromatin dynamics and the trans-histone H3 methylation. Epigenetics 2010, 5(6): 460–468.

18. Bonnet J, Devys D, Tora L. Histone H2B ubiquitination: signaling not scrapping. Drug Discov Today Technol 2014, 12: e19–27.

19. Bonnet J, Wang CY, Baptista T, Vincent SD, Hsiao WC, Stierle M, et al. The SAGA coactivator complex acts on the whole transcribed genome and is required for RNA polymerase II transcription. Gene Dev 2014, 28(18): 1999–2012.

20. Fuchs G, Hollander D, Voichek Y, Ast G, Oren M. Cotranscriptional histone H2B monoubiquitylation is tightly coupled with RNA polymerase II elongation rate. Genome research 2014, 24(10): 1572–1583.

21. Jung I, Kim SK, Kim M, Han YM, Kim YS, Kim D, et al. H2B monoubiquitylation is a 5’-enriched active transcription mark and correlates with exon-intron structure in human cells. Genome Res 2012, 22(6): 1026–1035.

22. Briggs SD, Xiao T, Sun ZW, Caldwell JA, Shabanowitz J, Hunt DF, et al. Gene silencing: trans-histone regulatory pathway in chromatin. Nature 2002, 418(6897): 498.

23. Dover J, Schneider J, Tawiah-Boateng MA, Wood A, Dean K, Johnston M, et al. Methylation of histone H3 by COMPASS requires ubiquitination of histone H2B by Rad6. J Biol Chem 2002, 277(32): 28368–28371.

24. Ng HH, Xu RM, Zhang Y, Struhl K. Ubiquitination of histone H2B by Rad6 is required for efficient Dot1-mediated methylation of histone H3 lysine 79. J Biol Chem 2002, 277(38): 34655–34657.

25. Sun ZW, Allis CD. Ubiquitination of histone H2B regulates H3 methylation and gene silencing in yeast. Nature 2002, 418(6893): 104–108.

26. Lee JS, Shukla A, Schneider J, Swanson SK, Washburn MP, Florens L, et al. Histone crosstalk between H2B monoubiquitination and H3 methylation mediated by COMPASS. Cell 2007, 131(6): 1084–1096.

27. Kim J, Kim JA, McGinty RK, Nguyen UT, Muir TW, Allis CD, et al. The n-SET domain of Set1 regulates H2B ubiquitylation-dependent H3K4 methylation. Mol Cell 2013, 49(6): 1121–1133.

28. Daniel JA, Torok MS, Sun ZW, Schieltz D, Allis CD, Yates JR, 3rd, et al. Deubiquitination of histone H2B by a yeast acetyltransferase complex regulates transcription. J Biol Chem 2004, 279(3): 1867–1871.

29. Henry KW, Wyce A, Lo WS, Duggan LJ, Emre NC, Kao CF, et al. Transcriptional activation via sequential histone H2B ubiquitylation and deubiquitylation, mediated by SAGA-associated Ubp8. Genes Dev 2003, 17(21): 2648–2663.

30. Zhao Y, Lang G, Ito S, Bonnet J, Metzger E, Sawatsubashi S, et al. A TFTC/STAGA module mediates histone H2A and H2B deubiquitination, coactivates nuclear receptors, and counteracts heterochromatin silencing. Mol Cell 2008, 29(1): 92–101.

31. Zhang XY, Varthi M, Sykes SM, Phillips C, Warzecha C, Zhu W, et al. The putative cancer stem cell marker USP22 is a subunit of the human SAGA complex required for activated transcription and cell-cycle progression. Mol Cell 2008, 29(1): 102–111.

32. Lang G, Bonnet J, Umlauf D, Karmodiya K, Koffler J, Stierle M, et al. The tightly controlled deubiquitination activity of the human SAGA complex differentially modifies distinct gene regulatory elements. Molecular and cellular biology 2011, 31(18): 3734–3744.

33. Morgan MT, Wolberger C. Recognition of ubiquitinated nucleosomes. Curr Opin Struct Biol 2017, 42: 75–82.

34. Atanassov BS, Mohan RD, Lan X, Kuang X, Lu Y, Lin K, et al. ATXN7L3 and ENY2 Coordinate Activity of Multiple H2B Deubiquitinases Important for Cellular Proliferation and Tumor Growth. Mol Cell 2016, 62(4): 558–571.

35. Bonnet J, Romier C, Tora L, Devys D. Zinc-finger UBPs: regulators of deubiquitylation. Trends Biochem Sci 2008, 33(8): 369–375.

36. Atanassov BS, Evrard YA, Multani AS, Zhang Z, Tora L, Devys D, et al. Gcn5 and SAGA regulate shelterin protein turnover and telomere maintenance. Mol Cell 2009, 35(3): 352–364.

37. Gennaro VJ, Stanek TJ, Peck AR, Sun Y, Wang F, Qie S, et al. Control of CCND1 ubiquitylation by the catalytic SAGA subunit USP22 is essential for cell cycle progression through G1 in cancer cells. Proc Natl Acad Sci U S A 2018, 115(40): E9298–E9307.

38. Atanassov BS, Dent SY. USP22 regulates cell proliferation by deubiquitinating the transcriptional regulator FBP1. EMBO Rep 2011, 12(9): 924–930.

39. Armour SM, Bennett EJ, Braun CR, Zhang XY, McMahon SB, Gygi SP, et al. A high-confidence interaction map identifies SIRT1 as a mediator of acetylation of USP22 and the SAGA coactivator complex. Mol Cell Biol 2013, 33(8): 1487–1502.

40. Lin Z, Yang H, Kong Q, Li J, Lee SM, Gao B, et al. USP22 antagonizes p53 transcriptional activation by deubiquitinating Sirt1 to suppress cell apoptosis and is required for mouse embryonic development. Mol Cell 2012, 46(4): 484–494.

41. Kobayashi T, Iwamoto Y, Takashima K, Isomura A, Kosodo Y, Kawakami K, et al. Deubiquitinating enzymes regulate Hes1 stability and neuronal differentiation. FEBS J 2015, 282(13): 2411–2423.

42. Lambies G, Miceli M, Martinez-Guillamon C, Olivera-Salguero R, Pena R, Frias CP, et al. TGFbeta-Activated USP27X Deubiquitinase Regulates Cell Migration and Chemoresistance via Stabilization of Snail1. Cancer Res 2019, 79(1): 33–46.

43. Zhou Z, Zhang P, Hu X, Kim J, Yao F, Xiao Z, et al. USP51 promotes deubiquitination and stabilization of ZEB1. Am J Cancer Res 2017, 7(10): 2020–2031.

44. Koutelou E, Wang L, Schibler AC, Chao HP, Kuang X, Lin K, et al. USP22 controls multiple signaling pathways that are essential for vasculature formation in the mouse placenta. Development 2019, 146(4).

45. Kosinsky RL, Wegwitz F, Hellbach N, Dobbelstein M, Mansouri A, Vogel T, et al. Usp22 deficiency impairs intestinal epithelial lineage specification in vivo. Oncotarget 2015, 6(35): 37906–37918.

46. Tyanova S, Temu T, Sinitcyn P, Carlson A, Hein MY, Geiger T, et al. The Perseus computational platform for comprehensive analysis of (prote)omics data. Nat Methods 2016, 13(9): 731–740.

47. Dobin A, Davis CA, Schlesinger F, Drenkow J, Zaleski C, Jha S, et al. STAR: ultrafast universal RNA-seq aligner. Bioinformatics 2013, 29(1): 15–21.

48. Love MI, Huber W, Anders S. Moderated estimation of fold change and dispersion for RNA-seq data with DESeq2. Genome Biol 2014, 15(12): 550.

49. El-Saafin F, Curry C, Ye T, Garnier JM, Kolb-Cheynel I, Stierle M, et al. Homozygous TAF8 mutation in a patient with intellectual disability results in undetectable TAF8 protein, but preserved RNA polymerase II transcription. Hum Mol Genet 2018, 27(12): 2171–2186.

50. Gyenis A, Umlauf D, Ujfaludi Z, Boros I, Ye T, Tora L. UVB induces a genome-wide acting negative regulatory mechanism that operates at the level of transcription initiation in human cells. PLoS genetics 2014, 10(7): e1004483.

51. Chapman RD, Heidemann M, Albert TK, Mailhammer R, Flatley A, Meisterernst M, et al. Transcribing RNA polymerase II is phosphorylated at CTD residue serine-7. Science 2007, 318(5857): 1780–1782.

52. Anders S, Huber W. Differential expression analysis for sequence count data. Genome Biol 2010, 11(10): R106.

53. Ye T, Krebs AR, Choukrallah MA, Keime C, Plewniak F, Davidson I, et al. seqMINER: an integrated ChIP-seq data interpretation platform. Nucleic acids research 2011, 39(6): e35.

54. Rahl PB, Lin CY, Seila AC, Flynn RA, McCuine S, Burge CB, et al. c-Myc regulates transcriptional pause release. Cell 2010, 141(3): 432–445.

55. Martello G, Smith A. The nature of embryonic stem cells. Annu Rev Cell Dev Biol 2014, 30: 647–675.

56. Hutchins AP, Yang Z, Li Y, He F, Fu X, Wang X, et al. Models of global gene expression define major domains of cell type and tissue identity. Nucleic Acids Res 2017, 45(5): 2354–2367.

57. Adelman K, Lis JT. Promoter-proximal pausing of RNA polymerase II: emerging roles in metazoans. Nature reviews Genetics 2012, 13(10): 720–731.

58. Krebs AR, Imanci D, Hoerner L, Gaidatzis D, Burger L, Schubeler D. Genome-wide Single-Molecule Footprinting Reveals High RNA Polymerase II Turnover at Paused Promoters. Mol Cell 2017, 67(3): 411–422 e414.

59. Erickson B, Sheridan RM, Cortazar M, Bentley DL. Dynamic turnover of paused Pol II complexes at human promoters. Genes Dev 2018, 32(17-18): 1215–1225.

60. Harlen KM, Churchman LS. The code and beyond: transcription regulation by the RNA polymerase II carboxy-terminal domain. Nat Rev Mol Cell Biol 2017, 18(4): 263–273.

61. Chen FX, Woodfin AR, Gardini A, Rickels RA, Marshall SA, Smith ER, et al. PAF1, a Molecular Regulator of Promoter-Proximal Pausing by RNA Polymerase II. Cell 2015, 162(5): 1003–1015.

62. Vincent SD, Dunn NR, Hayashi S, Norris DP, Robertson EJ. Cell fate decisions within the mouse organizer are governed by graded Nodal signals. Genes Dev 2003, 17(13): 1646–1662.

63. Seruggia D, Oti M, Tripathi P, Canver MC, LeBlanc L, Di Giammartino DC, et al. TAF5L and TAF6L Maintain Self-Renewal of Embryonic Stem Cells via the MYC Regulatory Network. Mol Cell 2019, 74(6): 1148–1163 e1147.

64. Laribee RN, Fuchs SM, Strahl BD. H2B ubiquitylation in transcriptional control: a FACT-finding mission. Genes Dev 2007, 21(7): 737–743.

65. Baptista T, Devys D. Saccharomyces cerevisiae Metabolic Labeling with 4-thiouracil and the Quantification of Newly Synthesized mRNA As a Proxy for RNA Polymerase II Activity. J Vis Exp 2018(140).

66. Fleming AB, Kao CF, Hillyer C, Pikaart M, Osley MA. H2B ubiquitylation plays a role in nucleosome dynamics during transcription elongation. Mol Cell 2008, 31(1): 57–66.

67. Batta K, Zhang Z, Yen K, Goffman DB, Pugh BF. Genome-wide function of H2B ubiquitylation in promoter and genic regions. Genes Dev 2011, 25(21): 2254–2265.

68. Fierz B, Kilic S, Hieb AR, Luger K, Muir TW. Stability of nucleosomes containing homogenously ubiquitylated H2A and H2B prepared using semisynthesis. J Am Chem Soc 2012, 134(48): 19548–19551.

69. Acharya D, Hainer SJ, Yoon Y, Wang F, Bach I, Rivera-Perez JA, et al. KAT-Independent Gene Regulation by Tip60 Promotes ESC Self-Renewal but Not Pluripotency. Cell Rep 2017, 19(4): 671–679.

70. Dorighi KM, Swigut T, Henriques T, Bhanu NV, Scruggs BS, Nady N, et al. Mll3 and Mll4 Facilitate Enhancer RNA Synthesis and Transcription from Promoters Independently of H3K4 Monomethylation. Mol Cell 2017, 66(4): 568–576 e564.

71. Rickels R, Herz HM, Sze CC, Cao K, Morgan MA, Collings CK, et al. Histone H3K4 monomethylation catalyzed by Trr and mammalian COMPASS-like proteins at enhancers is dispensable for development and viability. Nat Genet 2017, 49(11): 1647–1653.

