## Supplementary Figures, Supplementary figure legends, and Supplementary Table Legends for "Histone H2Bub1 deubiquitylation is essential for mouse development, but does not regulate global RNA polymerase II transcription"

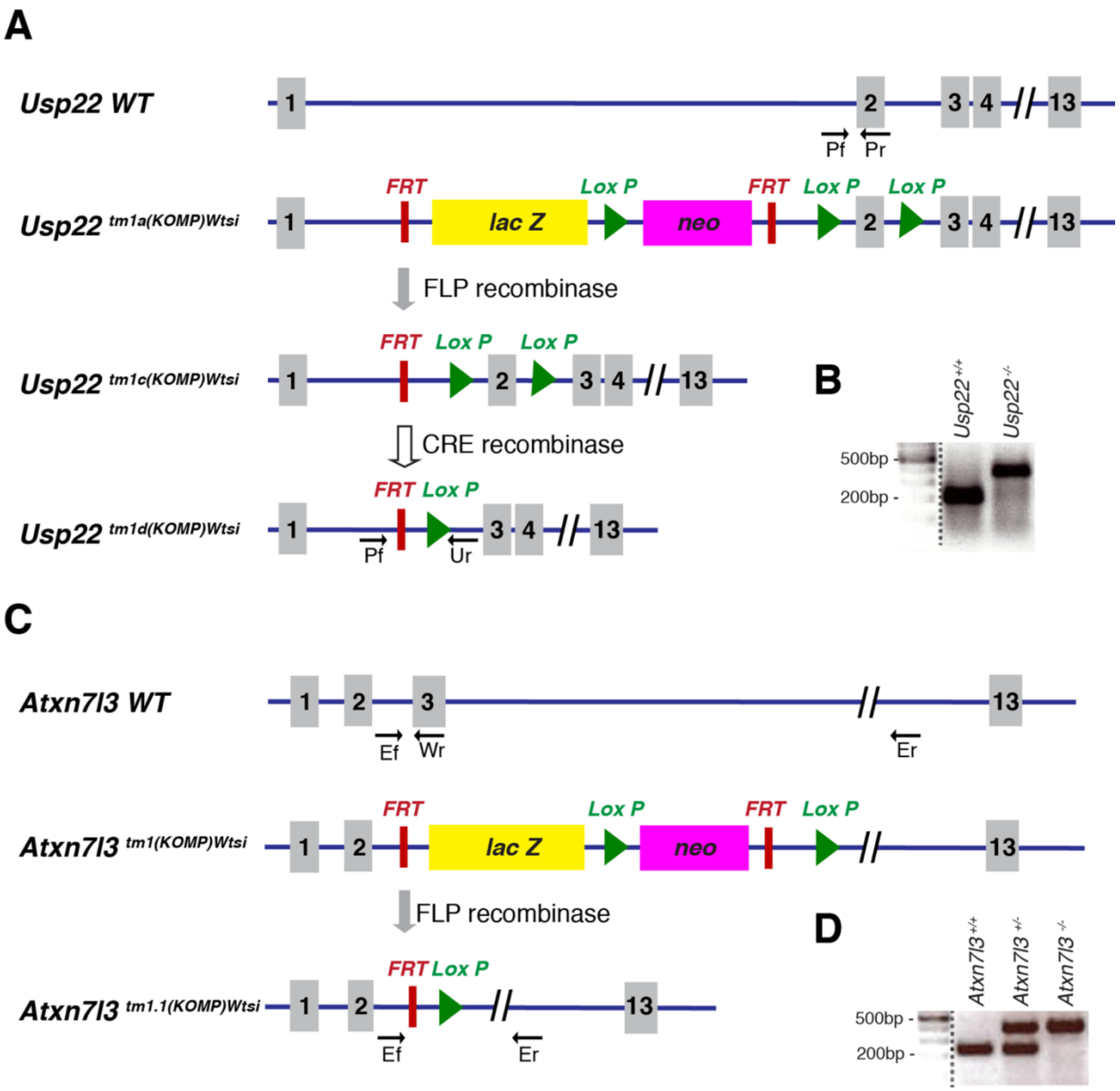


**Supplementary Figure 1: Deletion of the *Usp22* and *Atxn7l3* genes in mouse**

**A.** Generation of the *Usp22*$^{tm1d(KOMP)Wtsi}$ allele (*Usp22*$^{-}$) after FLP and CRE recombination of the *Usp22*$^{tm1a(KOMP)Wtsi}$ initial allele. The primers used for genotyping are indicated on the maps. **B.** PCR analysis of DNA samples using the Pf, Pr and Ur primers from *Usp22*$^{+/+}$ and *Usp22*$^{-/-}$ mice. The 199 and 362 bp bands correspond to the WT and null alleles, respectively. **C.** Generation of the *Atxn7l3*$^{tm1.1(KOMP)Wtsi}$ allele (*Atxn7l3*$^{-}$) after FLP recombination of the *Atxn7l3*$^{tm1(KOMP)Wtsi}$ initial allele. The primers used for genotyping are indicated on the maps. **D.** PCR analysis of DNA samples using the Ef, Er and Wr primers from *Atxn7l3*$^{+/+}$, *Atxn7l3*$^{+/-}$ and *Atxn7l3*$^{-/-}$ mice. The 215 and 318 bp bands correspond to the WT and null alleles, respectively.

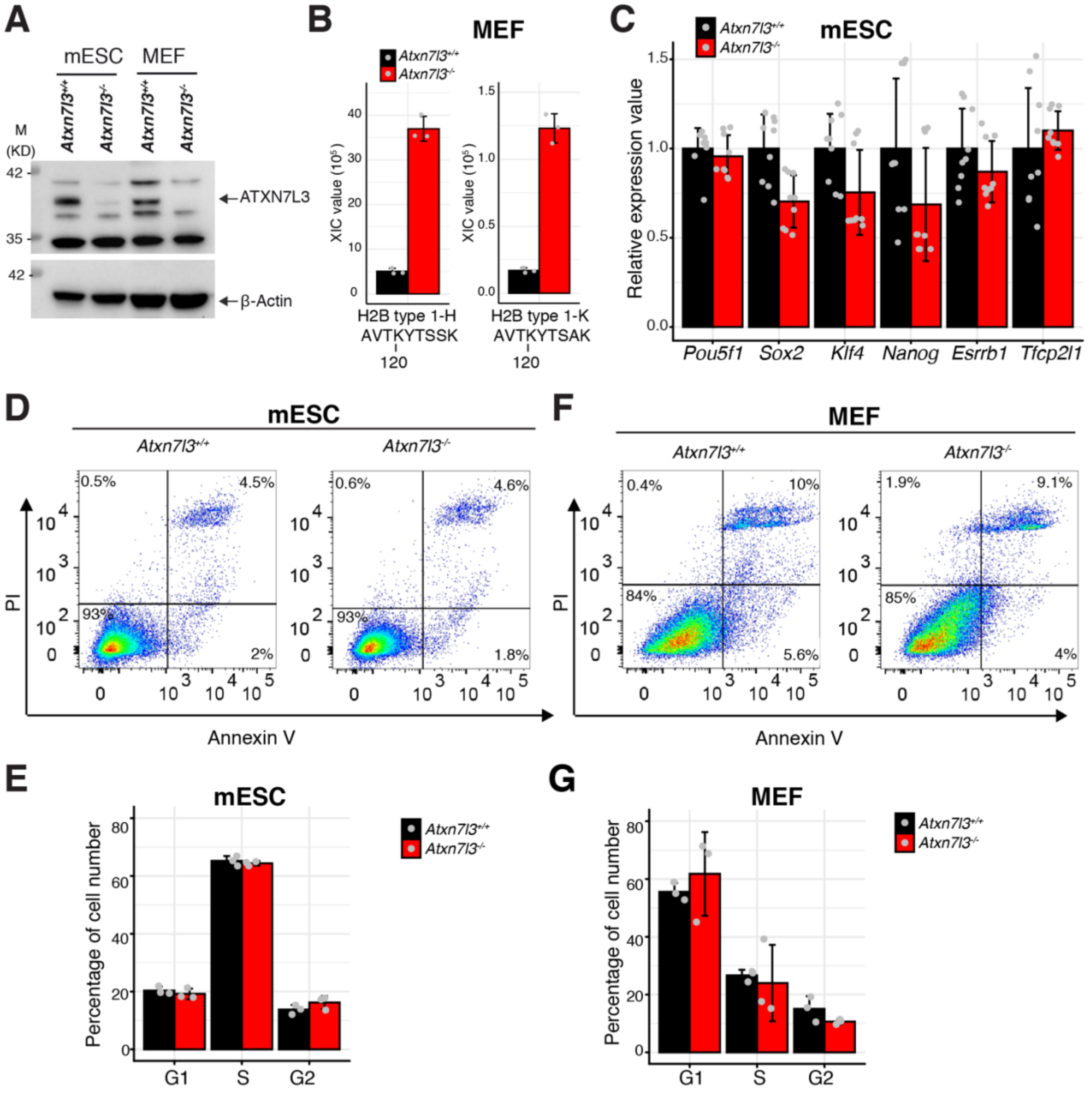


**Supplementary Figure 2:**

**A.** Western blot analyses of *Atxn7l3*$^{+/+}$ and *Atxn7l3*$^{-/-}$ mESC and MEF whole cell extracts using anti-ATXN7L3 and anti-β-Actin antibodies. M: molecular weight marker (in kDa). **B.** *Atxn7l3*$^{+/+}$ and *Atxn7l3*$^{-/-}$ MEF whole cell extracts were digested by endoproteinase Lys-C and trypsin, ubiquitinylated peptides were enriched with an anti-ubiquitin remnant motif (K-ε-GG) antibody, and analyzed by mass spectrometry. The identified monoubiquitylated histone H2B isoform peptides (the sequence of the two H2B peptides with the ubiquitylated lysine 120 are indicated) were determined in extracted-ion chromatogram (XIC) values and quantified (see Materials and Methods). **C.** RT-qPCR analysis of genes associated with pluripotency in *Atxn7l3*$^{+/+}$ (black) and *Atxn7l3*$^{-/-}$ (red) mESCs. Y axis indicates the relative mRNA expression to the *Pgk1* housekeeping gene. *Atxn7l3*$^{+/+}$ RNA expression level is normalized to 1. Error bars represent ±SD from three biological samples with three technical replicates (represented by grey dots) for each. **D** and **F.** Apoptosis measured by Annexin V and Propidium iodide (PI) staining and quantified by flow cytometry in mESC (D) and MEF (F) cells. **E** and **G.** Quantification of cell cycle phase distribution by flow cytometry from Propidium iodide (PI) treated *Atxn7l3*$^{+/+}$ (black) and *Atxn7l3*$^{-/-}$ (red) mESC (E) and MEF (G) cells. Error bars indicate ±SD based on three biological replicates (represented by grey dots).

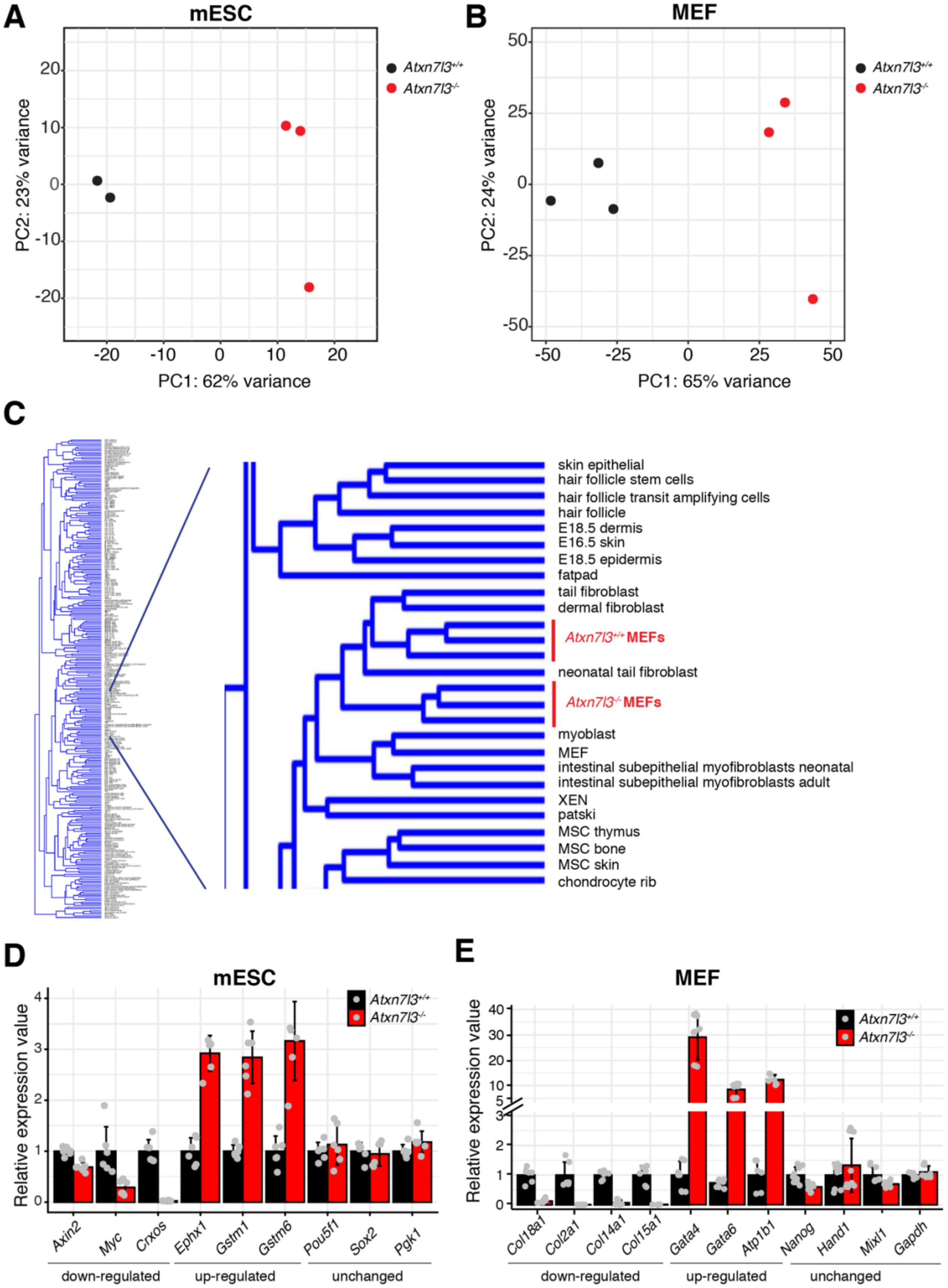


**Supplementary Figure 3**

**A-B.** Principal component analysis of control (black) and *Atxn7l3*$^{-/-}$ (red) RNA-seq data in mESCs (A) and MEFs (B). **C.** Hierarchical clustering of *Atxn7l3*$^{-/-}$ and control MEF RNA-seq data with 921 RNA-seq data from 272 distinct mouse cell types or tissues [1]. **D.** RT-qPCR analysis of up-regulated, down-regulated and unchanged genes from RNA-seq in mESC. Y axis indicates the relative mRNA expression to the *Pgk1* housekeeping gene in *Atxn7l3*$^{-/-}$ mESCs compared to WT controls. WT gene expression is normalized to 1. Error bars represent ±SD from two biological and three technical replicates (represented by grey dots). **E.** RT-qPCR analysis of up-regulated, down-regulated and unchanged genes from RNA-seq in *Atxn7l3*$^{-/-}$ MEFs compared to WT control. Y axis indicates the relative mRNA expression of a given gene normalized to the expression of *Pgk1 and Hsp90ab1* housekeeping genes. Error bars represent ±SD from two biological and three technical replicates (represented by grey dots).

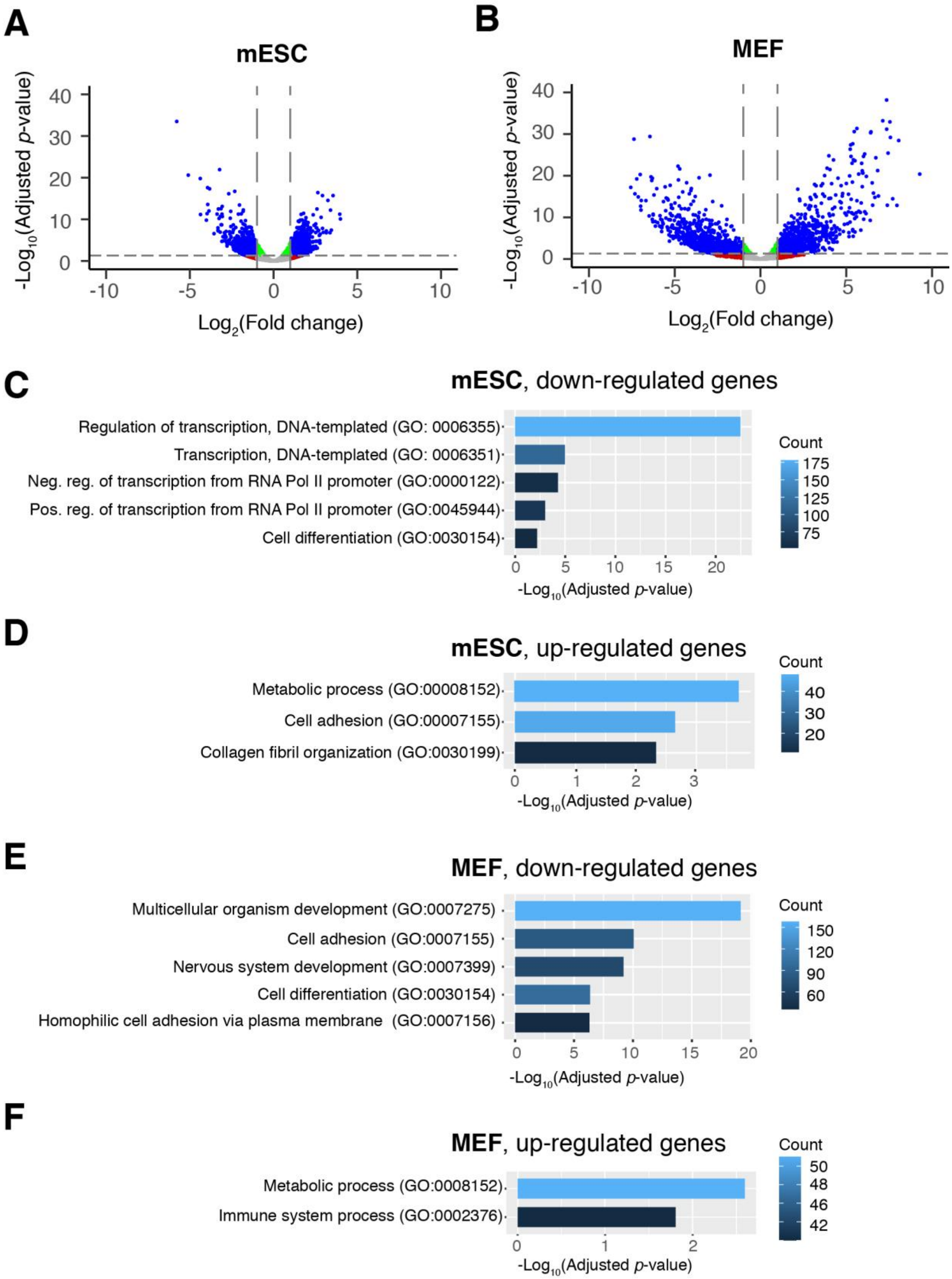


**Supplementary Figure 4**

**A-B.** Volcano plots comparing gene expression between *Atxn7l3*$^{-/-}$ and WT control mESCs (A) and MEFs (B). Blue dots correspond to significantly differentially expressed genes with adjusted *p*-values ≤ 0.05 and absolute $\log_2$(Fold change) ≥ 1. Green dots indicate genes with adjusted *p*-values < 0.05 and absolute $\log_2$(Fold change) < 1. Red dots indicate genes with adjusted *p*-values > 0.05 and absolute $\log_2$(Fold change) > 1. Grey dots indicate adjusted *p*-values > 0.05 and absolute $\log_2$(Fold change) < 1. **C-F.** Gene ontology (GO) analyses of differentially expressed genes both in *Atxn7l3*$^{-/-}$ mESCs, and *Atxn7l3*$^{-/-}$ MEFs versus WT controls. Results of gene ontology analyses carried out using DAVID bioinformatics resources 6.8 to identify differential gene-function categories (as indicated). Significantly enriched GO terms ($-\mathrm{Log}_{10}$ adjusted *p* value < 0.05) in biological processes are shown. The number of genes enriched in each GO category is also shown.

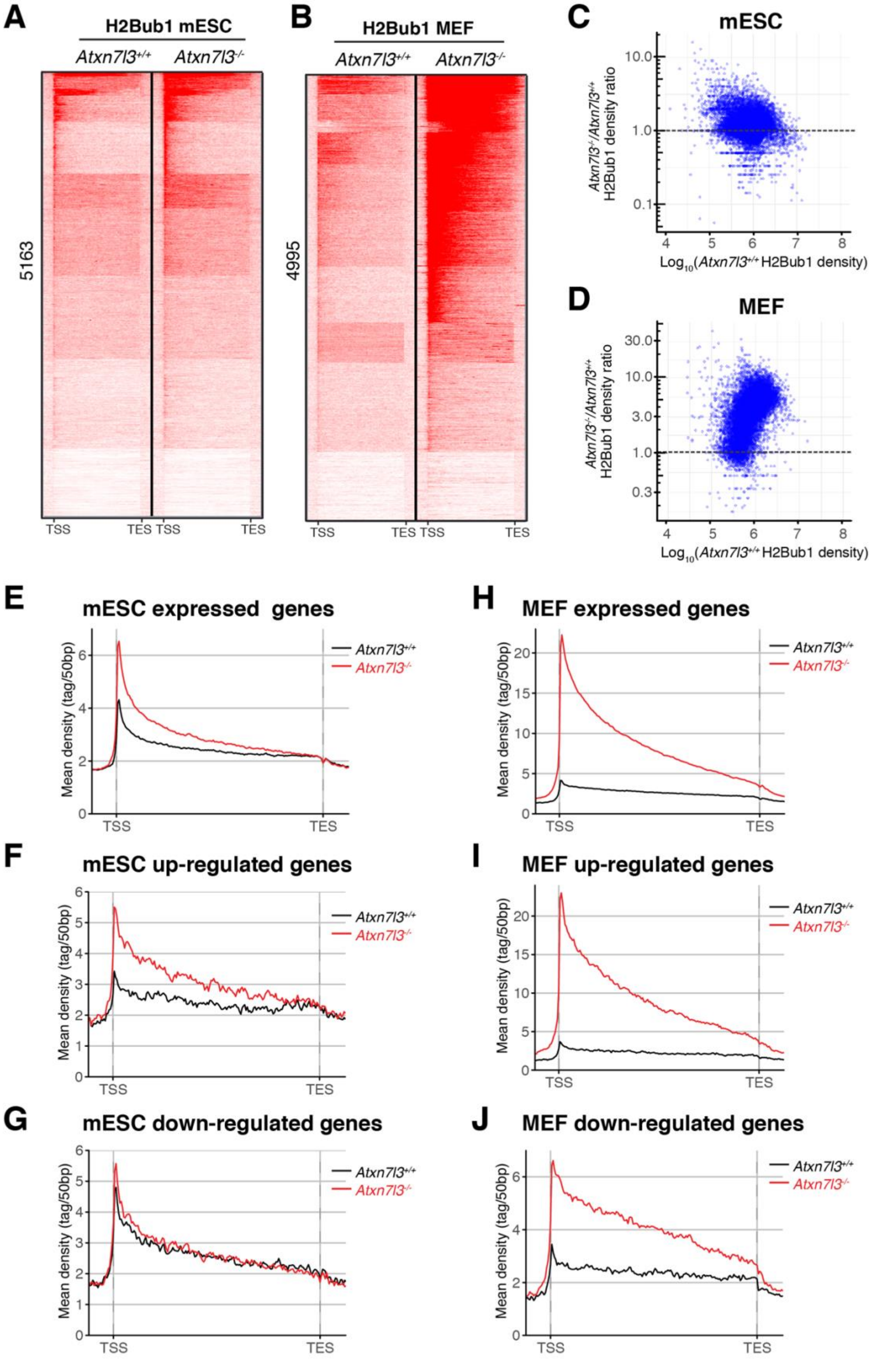
A
H2Bub1 mESC
Atxn7l3+/+ Atxn7l3-/-
5163
TSS TES TSS TES
B
H2Bub1 MEF
Atxn7l3+/+ Atxn7l3-/-
4995
TSS TES TSS TES
C
mESC
Atxn7l3-/-/Atxn7l3+/+ H2Bub1 density ratio
Log10(Atxn7l3+/+ H2Bub1 density)
D
MEF
Atxn7l3-/-/Atxn7l3+/+ H2Bub1 density ratio
Log10(Atxn7l3+/+ H2Bub1 density)
E
mESC expressed genes
F
mESC up-regulated genes
G
mESC down-regulated genes
H
MEF expressed genes
I
MEF up-regulated genes
J
MEF down-regulated genes
Mean density (tag/50bp)
Atxn7l3+/+
Atxn7l3-/-
TSS TES

**Supplementary Figure 5**

**A-B.** Heat maps showing the distribution of H2Bub1 on gene bodies of 5163 expressed transcripts in mESCs (A) and 4995 expressed transcripts in MEFs (B). Upstream of the TSSs -5 kb and downstream of the TESs +5 kb extensions were analyzed in the panels, and neighboring gene units which overlapped with these regions were removed from the expressed gene lists. **C-D.** Scatter plots representing H2Bub1 densities in WT cells relative to *Atxn7l3*$^{-/-}$ mESC (C) and MEF (D). From 16269 expressed transcripts in mESCs, 15467 expressed transcripts containing at least 1 read were selected (C; blue dots). From 15084 expressed transcripts in MEF cells, 14500 expressed transcripts containing at least 1 read were selected (D; blue dots). **E-G** Average metagene profiles showing H2Bub1 distribution on gene bodies of 5163 expressed, 301 up-regulated and 368 down-regulated transcripts in mESCs (from -5kb upstream of the TSS to +5 kb downstream of the TES, and as in (A and B) neighboring gene units, which overlapped with these extended gene regions were removed). **H-J** Average metagene profiles showing H2Bub1 distribution on gene bodies of 4995 expressed, 517 up-regulated and 624 down-regulated transcripts in MEFs (from -5kb upstream of the TSS to +5 kb downstream of the TES, and as in (A and B) neighboring gene units, which overlapped with these extended gene regions were removed).

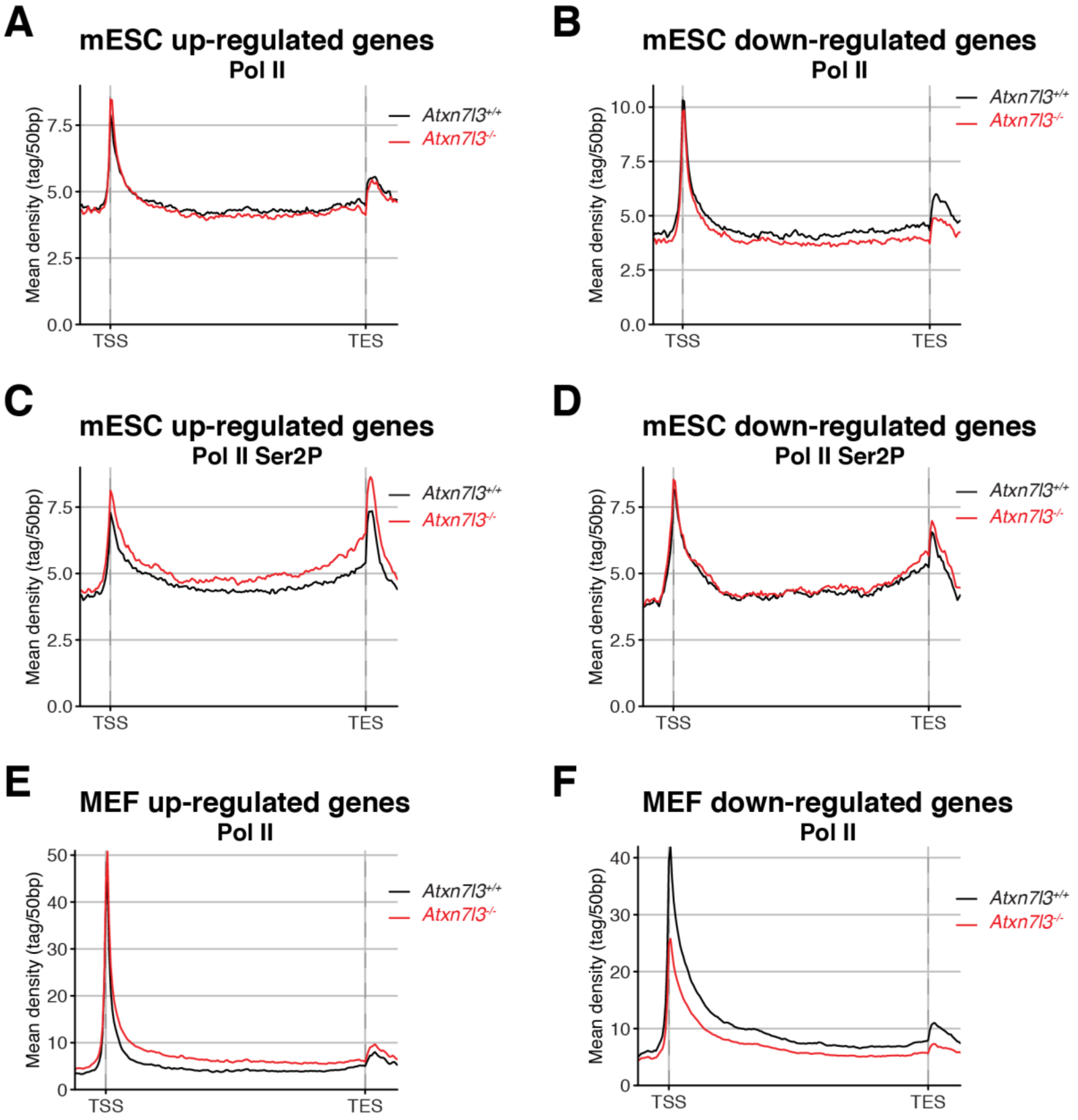


**Supplementary Figure 6**

**A-D.** Average metagene profiles showing Pol II (A and B) and Pol II-Ser2P (C and D) distribution on gene bodies of 1116 up-regulated and 810 down-regulated transcripts in mESCs (from -5kb upstream of the TSS to +5 kb downstream of the TES). **E-F.** Average metagene profiles showing Pol II distribution on gene bodies of 1185 up-regulated and 1555 down-regulated transcripts in MEFs (from -5kb upstream of the TSS to +5 kb downstream of the TES).

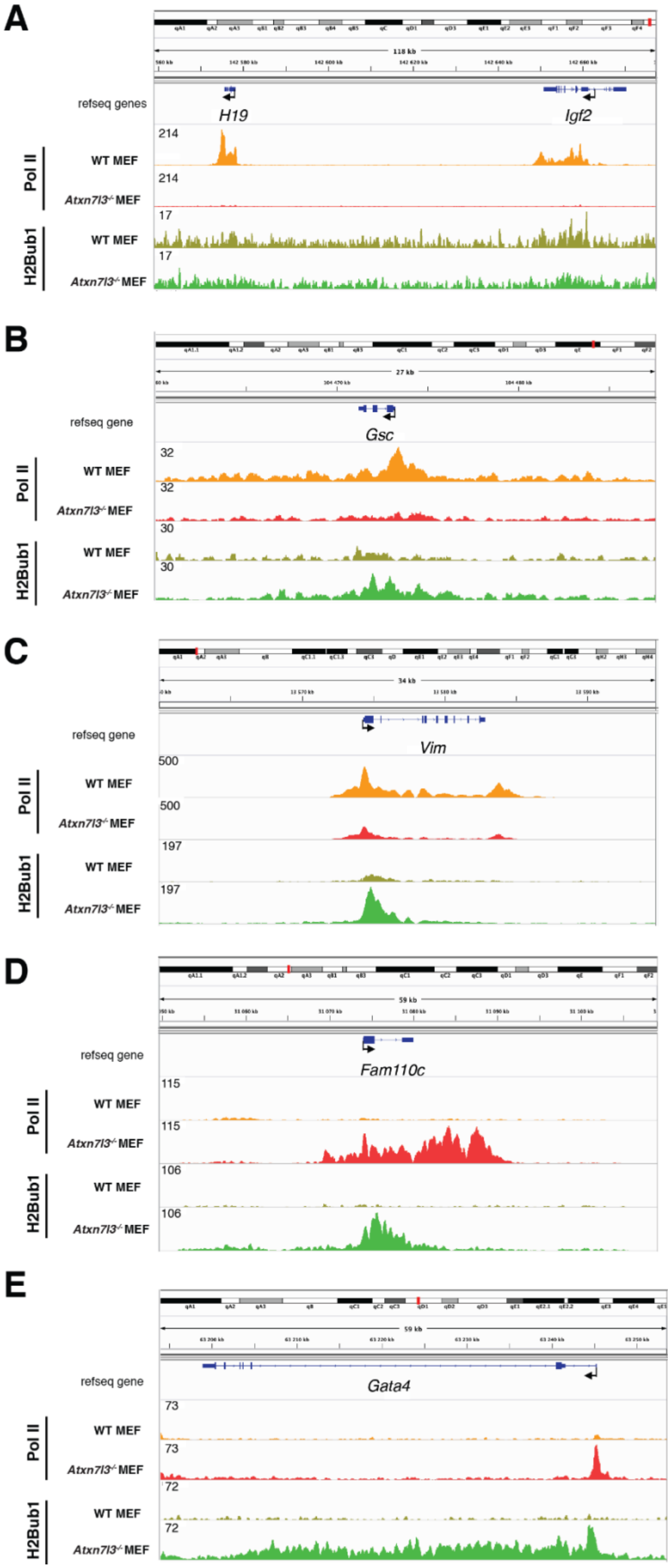


**Supplementary Figure 7**

**A-E.** H2Bub1 and Pol II binding profiles are shown at five selected genes in MEFs using the IGV genome browser. Direction of the transcription is indicated by arrows. Scaled tag densities for each gene are indicated on the left.

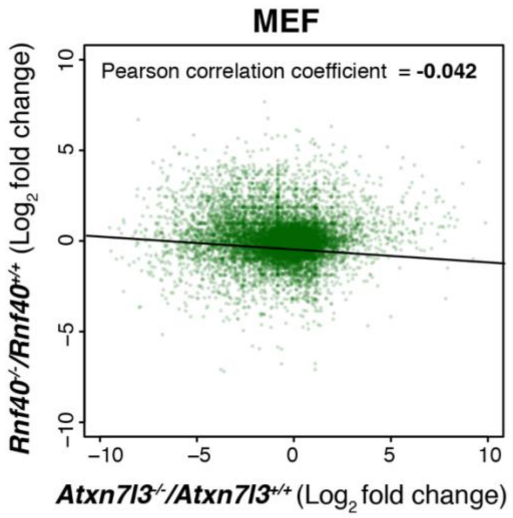


**Supplementary Figure 8**

Comparison of affected transcripts (in $\log_2$ fold change) in *Atxn7l3*$^{-/-}$/*Atxn7l3*$^{+/+}$ MEFs (our study) versus *Rnf40*$^{-/-}$/*Rnf40*$^{+/+}$ MEFs [2]. Each dot represents one gene. The linear regression line and the Pearson correlation coefficient are indicated.

**References**

1. Hutchins AP, Yang Z, Li Y, He F, Fu X, Wang X*, et al.* Models of global gene expression define major domains of cell type and tissue identity. *Nucleic Acids Res* 2017, **45**(5)**:** 2354-2367.

2. Xie W, Nagarajan S, Baumgart SJ, Kosinsky RL, Najafova Z, Kari V*, et al.* RNF40 regulates gene expression in an epigenetic context-dependent manner. *Genome Biol* 2017, **18**(1)**:** 32.

### Supplementary Tables

**Supplementary Table 1:** Primers used in the study

**Supplementary Table 2:** List of up- and down-regulated genes in mESC and MEFs

**Supplementary Table 3:** GO analysis Terms

**Supplementary Table 4:** Anti-H2Bub1 ChIP-seq counts in mESC and MEFs
